# TOR coordinates Cytokinin and Gibberellin signals mediating development and defense

**DOI:** 10.1101/2022.03.07.483332

**Authors:** Iftah Marash, Rupali Gupta, Gautam Anand, Meirav Leibman-Markus, Naomi Lindner, Alon Israeli, Dov Nir, Adi Avni, Maya Bar

## Abstract

Plants constantly perceive and process environmental signals and balance between the energetic demands of growth and defense. Growth arrest upon pathogen attack was previously suggested to result from a redirection of the plants’ metabolic resources towards the activation of plant defense. The energy sensor Target of Rapamycin (TOR) kinase is a conserved master coordinator of growth and development in all eukaryotes. Although TOR is positioned at the interface between development and defense, little is known about the mechanisms in which TOR may potentially regulate the relationship between these two modalities. The plant hormones cytokinin (CK) and gibberellin (GA) execute various aspects of plant development and defense. The ratio between CK and GA was reported to determine the outcome of developmental programs. Here, investigating the interplay between TOR-mediated development and TOR-mediated defense in tomato, we found that *TOR* silencing resulted in rescue of several different aberrant developmental phenotypes, demonstrating that TOR is required for the execution of developmental cues. In parallel, *TOR* inhibition enhanced immunity in genotypes with a low CK/GA ratio but not in genotypes with a high CK/GA ratio. TOR-inhibition mediated disease resistance was found to depend on developmental status, and was abolished in strongly morphogenetic leaves, while being strongest in mature, differentiated leaves. CK repressed TOR activity, suggesting that CK-mediated immunity may rely on TOR downregulation. At the same time, TOR activity was promoted by GA, and *TOR* silencing reduced GA sensitivity, indicating that GA signaling requires normal TOR activity. Our results demonstrate that TOR likely acts in concert with CK and GA signaling, executing signaling cues in both defense and development. Thus, differential regulation of TOR or TOR-mediated processes could regulate the required outcome of development-defense prioritization.

## Introduction

Plants have developed sophisticated strategies and molecular mechanisms to protect themselves against attacks by pathogens (Jiang et al., 2020). When subjected to biotic stresses, plants activate an array of cellular and molecular processes which include the production of defense proteins and metabolites. However, this activation of defense responses is often energy-demanding, and can suppresses plant growth by diverting energy and resources toward defense at the expense of growth, or by activation of conflicting pathways, or the sharing of components between immune and growth signaling (Eichmann & Schäfer, 2015). This is known as the ‘growth-defense tradeoff’, a phenomenon in which plants must constantly regulate and balance growth and defense in order to adapt to changes. It is now accepted that the tradeoff between growth and defense is carefully regulated by the plant, rather than a passive process in which energy diverted toward defense is simply not available for other needs (Karasov et al., 2017; Kliebenstein, 2016). Yet, our understanding of the mechanisms that enable plants to balance growth during biotic stress response is still limited.

In recent years, the conserved Target of Rapamycin (TOR) kinase has been established as a central eukaryotic regulatory hub, playing a role in the regulation of various cellular processes including metabolism, mRNA translation and transcription, cell division, rRNAs and ribosomal proteins synthesis, and autophagy (Dobrenel et al., 2016; Zhang et al., 2016). The TOR signaling pathway fine-tunes growth and development by coordinating nutrient availability, energy status and external cues. In plants, TOR signaling is particularly important for embryogenesis, meristem activation, leaf and root growth, senescence, and flowering (McCready et al., 2020). Under nutrient availability and when conditions are favorable for growth, the TOR signaling pathway is activated and developmental and anabolic processes are promoted while catabolic processes are repressed. When nutrients are limited or in the presence of environmental stresses, TOR is inactive and catabolic processes are promoted (Dobrenel et al., 2016; Saxton & Sabatini, 2017). Even though TOR signaling is involved in the regulation of multiple important signaling pathways, there is currently limited evidence of its crosstalk with the plant hormones GA and CK, and its possible involvement in CK-mediated immunity.

Recent studies suggest that TOR acts as a negative regulator of plant immunity, as it has been demonstrated to antagonize the defense hormones JA and SA in rice, suggesting that it may act as a switch between these modalities (De Vleesschauwer et al., 2018). Similarly, TOR was reported to negatively regulate JA biosynthesis and response in cotton (Song et al., 2017). Furthermore, mutants impaired in TOR complex and TOR-inhibited WT *Arabidopsis* plants were more resistant to *Fusarium* (Aznar et al., 2018), and in citrus spp., TOR inhibition was found to attenuate the growth of *Xanthomonas citri* (Soprano et al., 2018). In another study, *TOR* expression was downregulated upon NB-LRR activation. Suppression of *TOR* expression enhanced disease resistance, whereas *TOR* overexpression decreased it, suggesting that translational regulation executed by TOR plays an important role in the switch from growth to defense (Meteignier et al., 2018). TOR inhibition was also found to block growth and to activate the SA signaling pathway in *Arabidopsis* (Dong et al., 2015; Moreau et al., 2012). In agreement with this, we previously showed that TOR inhibition or *TOR* silencing promotes resistance against *Xanthomonas*, Tobacco mosaic virus (TMV), *Alternaria alternata* and *Botrytis cinerea* (*Bc*) in tomato and *N. benthamiana*, by SA-dependent activation of plant defense responses (Marash et al., 2022). Although the exact mechanism by which the inhibition of TOR primes resistance is not fully understood, it was suggested to selectively regulate translational control during plant immunity (Meteignier et al., 2018) and/or negatively regulate autophagy in plants, as was reported in yeast and mammals (Liu & Bassham, 2010).

Recent studies have shown that the TOR signaling pathway interacts with several plant hormones. TOR signaling interacts with the Brassinosteroid (BR) signaling pathway during hypocotyl elongation through the BZR1 transcription factor (Zhang et al., 2016), and activates Abscisic acid (ABA) receptors by phosphorylation (Wang et al., 2018). Additionally, TOR phosphorylates and stabilizes the Auxin (AUX) efflux facilitator PIN2, which affects the distribution gradient of PIN2 in *Arabidopsis* primary roots (Yuan et al., 2020). TOR monitors the level of sugar in meristematic regions and halts growth when the sugar level is low, blocking hormone signals that normally promote growth (Xiong et al., 2013). As plant growth rate is dictated by hormones, it seems that energy status and growth are integrated through the activity of TOR (Monson et al., 2022). Furthermore, TOR inhibition was shown to alter the expression of hundreds of genes, including genes that are linked to plant hormone signaling networks. When TOR is inhibited, the expression of genes involved in the signaling of growth hormones (AUX, GA, BR and CK) is repressed, while the expression of stress / growth inhibiting hormones (ABA, JA and SA) is upregulated (Dong et al., 2015). Although these findings demonstrated the existence of a relationship between TOR, GA, and CK signaling, the role of TOR in GA and CK-mediated immunity remains unclear.

Cytokinin (CK) is a plant hormone that regulates many aspects of plant growth and development including cell division, leaf senescence, apical dominance, vascular differentiation, chloroplast biogenesis, root development and stress responses (Zürcher & Müller, 2016). Previous studies have shown that CKs have a role in plant response to biotic stresses in tobacco (Großkinsky et al., 2011), Rice (Jiang et al., 2013) and tomato (Gupta et al., 2020). Several studies have reported that CKs promote resistance through the SA signaling pathway (Choi et al., 2010; Naseem et al., 2012). In tomato, we have previously shown that CK-deficiency results in higher susceptibility to the fungi *Botrytis cinerea* (*Bc*) and *Oidium neolycopersici* (*On*), while high endogenous CK content, as well as external application of CK, confer increased resistance against these fungi, in a SA-defendant manner (Gupta et al., 2020). Moreover, we have shown that CKs directly inhibit the growth, development, and virulence of fungal pathogens (Gupta et al., 2021), and that CKs improve *Xanthomonas campestris pv. Vesicatoria* (*Xcv*) and *Pseudomonas syringae* pv. *tomato Pst* disease outcome in tomato (Gupta et al., 2021).

Gibberellins (GAs) are growth-promoting phytohormones that play critical roles throughout the plant’s life cycle, including stem elongation, germination, leaf expansion, flowering, and fruit development (Davière & Achard, 2013). GAs regulate growth by destabilizing DELLA, a class of nuclear growth-repressing proteins that act as key regulators of the GA signaling and inhibit GA responses by interaction with multiple transcription factors (Locascio et al., 2013). Binding of GA to its receptor GA INSENSITIVE DWARF (GID1) results in degradation of DELLA and activation of responsive genes in the GA signaling pathway (Harberd et al., 2009; Hauvermale et al., 2012). GAs regulate plant growth in response to environmental changes as well as nutrient availability (Colebrook et al., 2014). Previous works demonstrated that GAs act as negative regulators of JA signaling (Campos et al., 2016; Major et al., 2020).

The opposing effects of GA and CK on many aspects of plant growth and development, such as shoot apical meristem formation, shoot and root elongation, and cell differentiation, often lead to their perception as antagonists (Ezura & Harberd, 1995; Jasinski et al., 2005). For instance, treatment with CK reduces GA activity by downregulation of GA biosynthesis genes and upregulation of two DELLA genes, *GAI* and *RGA* (Brenner et al., 2005). In addition, Greenboim-Wainberg et al., 2005 have shown that GA inhibits CK responses in Arabidopsis. The balance between CK and GA is maintained by three main proteins: KNOX, SPY and SEC. KNOX proteins induce CK biosynthesis while inhibiting GA biosynthesis and promoting GA deactivation. On the other hand, SPY and SEC repress GA signal and promote CK signals (Weiss & Ori, 2007). CK and GA have development-reciprocal relations, in which CK inhibits GA biosynthesis and promotes its deactivation by DELLA, and GA inhibits CK response. Normal shoot apical meristem function requires high CK and low GA signal, whereas later developmental stages require the opposite: low CK and high GA signals (Weiss & Ori, 2007). Considering that TOR and CK were both implicated in SA-dependent plant responses to pathogens, it appears possible that TOR and CK may interact or share similar defense pathways.

Here, we assessed the involvement of TOR in the mediation of GA and CK signals in both immunity and development. By inhibiting or down-regulating TOR, we observed partial rescue of abnormal development and defense phenotypes caused by imbalanced CK or GA levels. Our findings suggest that TOR plays a role in the mediation of developmental and defense signals originating from the balance between CK and GA.

## Results

### Leaf age and CK/GA balance affect plant immunity

Age has been previously linked to alterations in disease susceptibility (Goss & Bergelson, 2006). To examine *Botrytis cinerea (Bc)* susceptibility in the context of “leaf developmental age”, we first compared *Bc* sensitivity across different leaves on the same plants. We found that younger tomato leaves (L8) are significantly more resistant to *Bc* than older leaves (L5 and L3) **(Figure 1A)**. It is well established that the plant leaf developmental program is primarily influenced by the CK/GA balance (Fleishon et al., 2011; Hay et al., 2004; Shani et al., 2006). Young leaves undergoing morphogenesis are usually high in CK and low in GA, while differentiated organs have an opposite CK/GA balance (Israeli et al., 2021; Shwartz et al., 2016). Plant with high endogenous levels of CK or increased CK signaling age more slowly, have delayed reproduction and senescence, and retain “juvenile” developmental characteristics for longer periods (Gan & Amasino, 1995; Hwang et al., 2006, 2012; Riefler et al., 2006). We therefore hypothesized that the reduction in *Bc* disease severity in young tissues could relate to the CK/GA ratio. To test this hypothesis and to better understand the mechanisms that may confer resistance to plants with high CK levels, we investigated *Bc* susceptibility in leaves of different developmental stages in genotypes with altered CK/GA ratios. We used the following genotypes, all in the M82 background: *pBLS>>IPT7*, which contains elevated endogenous levels of CK (Shani et al., 2010) referred to hereinafter as “*pBLS>>IPT* “or “*IPT*”; *clausa*, which has increased CK sensitivity coupled with decreased CK content (Bar et al., 2016), referred to hereinafter as “*clausa*” or “*clau*”; *pFIL>>CKX3*, which has reduced CK levels (Shani et al., 2010; Shwartz et al., 2016), referred to hereinafter as “*pFIL>>CKX*” or “*CKX*”; *pFIL>>GFP-PROΔ17*, which has low GA signaling, referred to hereinafter as “*pFIL>> PROΔ17*” or “*PROΔ17*” (Nir et al., 2017); *ga20ox3*, which is predicted to have reduced GA levels; and *procera^ΔGRAS^*, referred to hereinafter as “*procera*” or “*pro*”, which has increased GA signaling (Livne et al., 2015). All the genotypes used in this study are detailed and justified in **Table 1** in the materials section.

**Figure 1:**
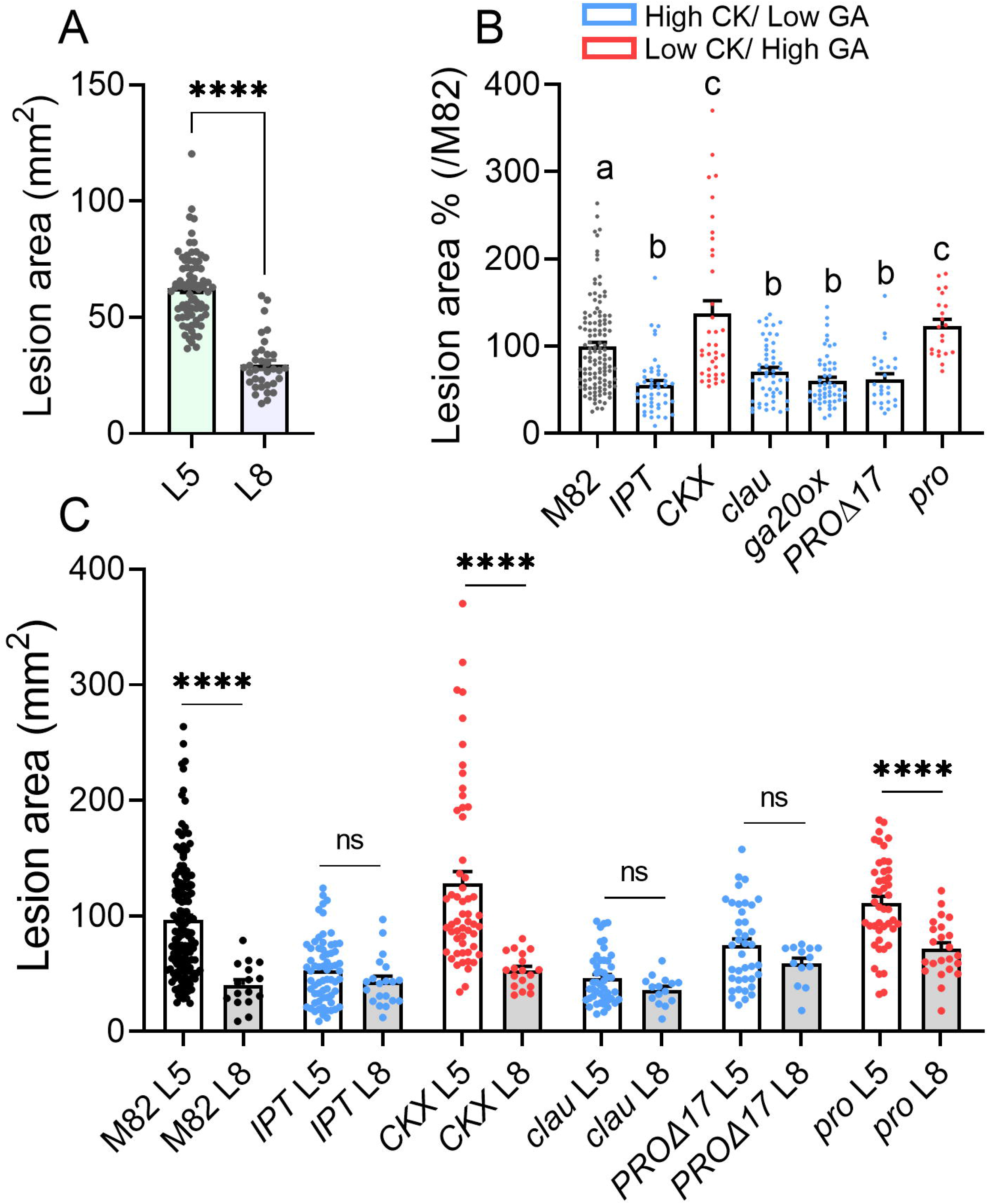
Disease susceptibility depends on leaf age and hormonal balance. **A**: Different leaves as indicated from *S. lycopersicum* cv. M82 5 week old plants were infected with *B. cinerea*. Asterisks denote staistical significance among indicated samples in Welch’s t-test N>35, ****p<0.0001. **B:** The fifth leaf of *S. lycopersicum* 5 week old plants of altered CK/GA genotypes: increased CK content *pBLS>>IPT7* (“*IPT*”), decreased CK content *pFIL>>CKX3* (“*CKX*”), increased CK sensitivity and decreased GA sensitivity *clausa* mutant (“*clau*”), decreased GA content mutant (“*ga20ox*”), decreased GA signaling *pFIL>>proΔ17* (“*proΔ17*”), increased GA signaling *procera* (“*pro*”) and their WT background M82, were infected with *B. cinerea* mycelia from a 72h old-culture. Different letters indicate staistically significant differences among indicated samples in Welch’s ANOVA with Dunnett’s post hoc test, N>20, p<0.018. **C**: Different leaves as indicated from *S. lycopersicum* 5 week old plants of altered CK/GA genotypes: increased CK content *pBLS>>IPT7* (“*IPT*”), decreased CK content *pFIL>>CKX3* (“*CKX*”), increased CK sensitivity and decreased GA sensitivity *clausa* mutant (“*clau*”), decreased GA signaling *pFIL>>proΔ17* (“*proΔ17*”), increased GA signaling *procera* (“*pro*”) and their WT background M82, were infected with *B. cinerea* mycelia from a 72h old-culture. Asterisks denote staistical significance among indicated samples in Welch’s ANOVA with Dunnett’s post hoc test, N>13, ****p<0.0001, ns-non significant. Bars represent Mean ±SEM, all points shown.

**Table 1:** Plant genotypes used in this study.

| Genotype name | Source | Phenotype | References |
| --- | --- | --- | --- |
| *S. lycopersicum* cultivar M82 |  | WT |  |
| *ga20ox3* null mutant<br>Cas9 Knock out of the GA biosynthesis enzyme GA20OX3. M82 background line. | Prof. Naomi Ori | Short stature; increased leaf complexity. | This work |
| *pTCSv2::3xVENUS*<br>Overexpression of VENUS driven by the synthetic two-component signaling sensor pTCSv2. M82 background line. | Bar lab | WT | (Bar et al., 2016; Steiner et al., 2020) |
| *pBLS>>IPT7 ("IPT")*: Transgenic overexpression of IsoPentenyl Transferase7 from the leaf *BLS* (Lifschitz et al., 2006) promoter. Elevated endogenous leaf levels of CK. M82 background line. | Prof. Naomi Ori | Highly complex rugose leaves; short stature; hirsutism; disease<sup>n,b</sup> resistance. | (Bar et al., 2016; Gupta et al., 2020, 2022; Gupta, Leibman-Markus, et al., 2021; Shani et al., 2010) |
| *pFIL>>CKX3 ("CKX")*: Transgenic overexpression of Cytokinin Oxidase3 from the leaf *FIL* (Bonaccorso et al., 2012) promoter. Decreased endogenous leaf CK content. M82 background line. | Prof. Naomi Ori | Highly simple thin leaves; lack of hairy trichomes; disease<sup>n,b</sup> susceptibility. | (Bar et al., 2016; Gupta et al., 2020, 2022; Shani et al., 2010) |
| *clausa ("clau")*: Recessive MYB TF mutant. High CK sensitivity coupled | Prof. Naomi Ori | Highly complex rugose leaves; | (Bar et al., 2016; Gupta et al., 2020, |
| with low CK content; low GA sensitivity coupled with increased amounts of pre-active GAs. M82 background line. |  | hirsutism; disease resistance <sup>n,b</sup>. | 2022; Shani et al., 2010) |
| *pFIL>>GFP-PROΔ17*: Transgenic overexpression of a mutated, un-processed version of the DELLA TF *Procera* from the leaf *FIL* promoter. Low GA signal. M82 background line. | Prof. David Weiss | Complex leaves. Disease resistance <sup>n,b</sup> (this work). | (Israeli et al., 2021; Nir et al., 2017) |
| *procera*: Recessive mutant in the DELLA TF *Procera*. ΔGRAS allele. High GA signal. M82 background line. | Prof. David Weiss | Simple leaves; tall stature. Disease susceptibility <sup>n</sup>/ resistance <sup>b</sup> (this work; resistance <sup>b</sup> previously reported in *Arabidopsis*). | (Livne et al., 2015; Navarro et al., 2008) |

<sup>n</sup> necrotrophic
<sup>b</sup> hemi/biotrophic

As shown in **Figure 1B**, when compared with the background line M82, genotypes with high CK/GA ratio (*IPT*, *clausa*, *ga20ox*, *PROΔ17*) exhibit significantly higher resistant to *Bc*, whereas genotypes with low CK/GA ratio (*CKX*, *pro*) are significantly more sensitive to *Bc*. This finding is in agreement with our and other previous studies (Bari & Jones, 2009; De Bruyne et al., 2014; Gupta et al., 2020; Wang et al., 2013). To examine the connection between the CK/GA ratio and leaf developmental age-related resistance, we examined the age-related *Bc* sensitivity of genotypes with altered CK/GA ratios. We found that leaf developmental age-related resistance was preserved when the CK/GA ratio was low (*CKX*, *pro*), but abolished when the CK/GA ratio was high (*IPT*, *clausa*, *PROΔ17*) (**Figure 1C**), indicating that CK-mediated resistance supersedes and/or is the same as leaf developmental age-related resistance.

### TOR inhibition mediates disease resistance and immunity effected by the CK/GA ratio

In our previous work, we showed found that similar to CK-mediated resistance, downregulation of TOR promotes immunity through the SA pathway (Marash et al., 2022). As it emerged from our results that the CK/GA ratio affects not only development but also disease resistance, we next turned to examine the connection between TOR and the CK/GA ratio in the context of immunity. We used Torin2, a potent ATP-competitive inhibitor of TOR activity in plants (Shi et al., 2018; Xiong et al., 2013; Ye et al., 2022) to inhibit TOR, and found that it reduced *Bc* disease symptoms in the M82 background and in genotypes with low CK/GA ratio (*CKX, pro*). However, we did not observe any additional decrease in *Bc* disease level when TOR was inhibited in genotypes with high CK/GA ratios (*IPT, clau, ga20ox, PROΔ17*) **(Figure 2A)**. We confirmed these results using another TOR inhibitor, WYE132 (**Figure S1**). We also silenced the expression of the tomato *SlTOR* gene by virus-induced gene silencing (VIGS), which reduces *SlTOR* transcription level by 50% in *TRV2:SlTOR* leaves (Marash et al., 2022), and observed similar results (**Figure S2**). This suggests that TOR might be required to transmit the output of the GA/CK ratio in defense.

**Figure 2:**
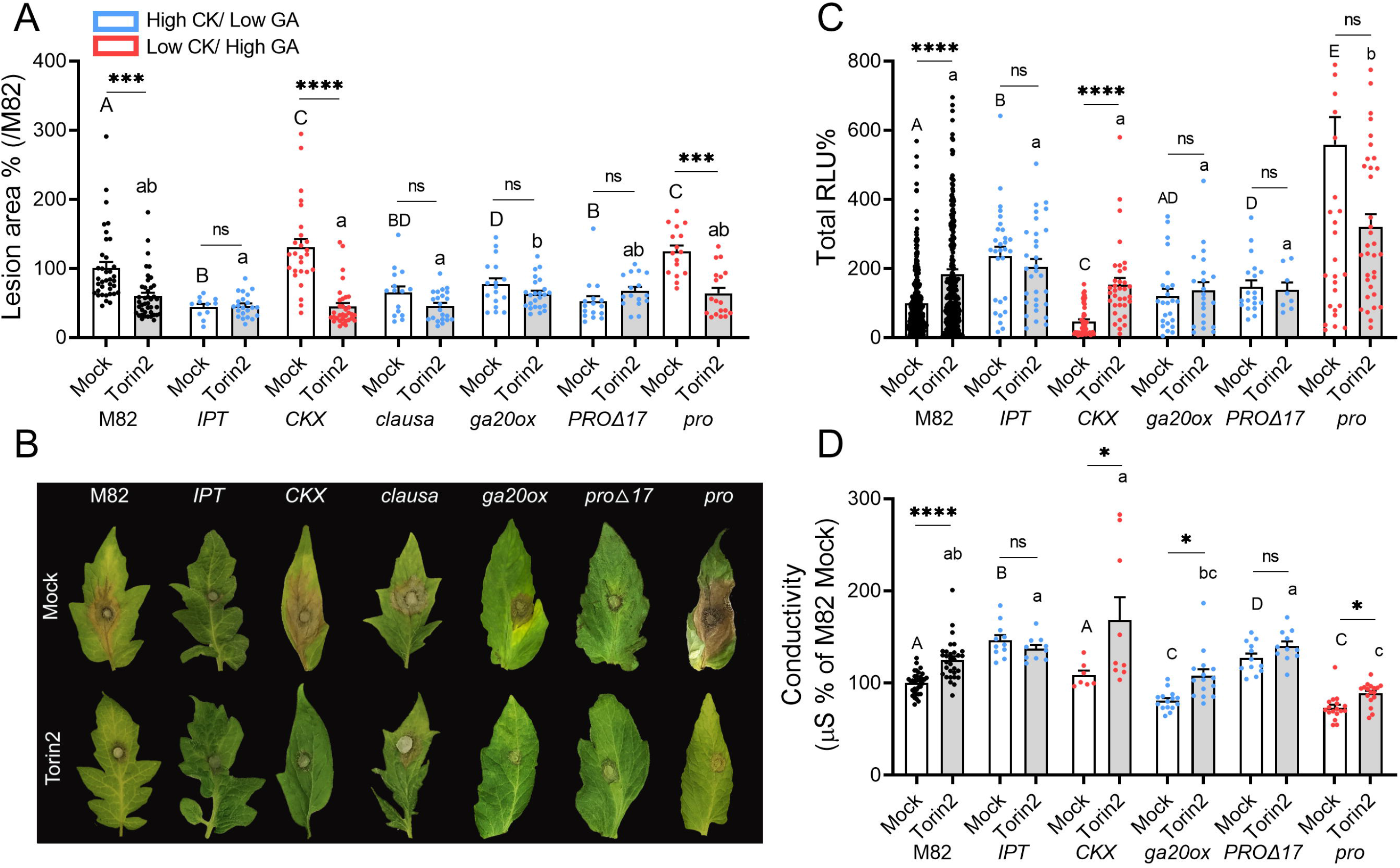
TOR inhibition mediated disease resistance depends on the CK/GA balance. *S. lycopersicum* plants of altered CK/GA genotypes: increased CK content *pBLS>>IPT7* (“*IPT*”), decreased CK content *pFIL>>CKX3* (“*CKX*”), increased CK sensitivity and decreased GA sensitivity *clausa* mutant (“*clau*sa”), decreased GA content mutant (“*ga20ox*”), decreased GA signaling *pFIL>>proΔ17* (“*proΔ17*”), increased GA signaling *procera* (“*pro*”) and their WT background M82, were treated with Mock (1:5000 DMSO in DDW), or 2 µM Torin2. Plants were challenged with *B. cinerea (Bc)* mycelia from a 72h old-culture 24 h after treatment. **A**: *Bc* necrotic lesion size. Asterisks indicate statistically significant disease reduction upon Torin2 treatment when compared with Mock treatment. Different letters indicate statistically significant differences among samples, upper case letters for Mock treated genotypes and lower case letters for samples treated with Torin2, in a one-way ANOVA with a Tukey post hoc test, N>12, p<0.044 (***p<0.001, ****p<0.0001, ns-non significant). Experiments were repeated 6 independent times. **B**: Representative *Bc* infected leaf images. **C**: Plants were challenged with the immunity elicitor flg-22 (1 µM) 24 h after Torin2 treatment. ROS production was measured immediately after flg-22 application every three minutes, using the HRP-luminol method, and expressed as Relative Luminescent Units (RLU). Average total RLU per treatment, expressed as % of M82 control, is plotted. **D**: Conductivity as a results of wounding was measured 24 h after Torin2 treatment. Bars represent mean ±SEM, all points shown. Experiments were repeated 3 independent times. **C, D**: Asterisks indicate statistically significant increases in ROS production or conductivity upon Torin2 treatment when compared with Mock treatment. Different letters indicate statistically significant differences among samples, upper case letters for Mock treated genotypes and lower case letters for samples treated with Torin2 in Kruskal-Wallis ANOVA with Dunn’s post-hoc test, N>20, p<0.0001.

As differences in *Bc*-sensitivity could be due to changes in cellular immunity, we examined the effect of TOR inhibition on defense responses in genotypes with an altered CK/GA ratio. In mock treated samples, consistent with our previous report characterizing CK-mediated immunity (Gupta et al., 2020), flg-22 elicited ROS levels were higher in *IPT*, and lower in *CKX* plants (**Figures 2C, S3**). Flg-22 elicited ROS production was enhanced in the mock samples of the high CK/GA ratio genotype *PROΔ17*, and interestingly, in the low CK/GA ratio genotype *pro* (**Figures 2C, S3**). This increased ROS production in *pro* could stem from higher levels of SA, similar to the increased SA levels observed in the quadruple-DELLA mutant, which lacks four out of the five DELLA proteins in *Arabidopsis* (Navarro et al., 2008). Torin2 treatment led to increased ROS production in M82 plants as previously reported (Marash et al., 2022) and in the low CK/GA ratio genotype *CKX* **(Figures 2C, S3)**. However, Torin2 did not affect any of the the high CK/GA ratio genotypes *IPT*, *ga20ox3*, or *PROΔ17* (**Figures 2C, S3**). We did not observe any significant change in the low CK/GA ratio genotype *pro*, again, possibly due to extremely high initial ROS levels. Similar results were obtained when quantifying ion leakage in response to Torin2, with increased in the background M82 line and in the low CK/GA ratio genotypes *CKX* and *pro*, but not in the high CK/GA genotypes, apart from *ga20ox*3 (**Figure 2D**).

We continued to assess the effect of TOR downregulation on disease resistance in lines with an altered CK/GA ratio by examining susceptibility to the hemibiotrophic bacterial pathogen *Xanthomonas campestris pv. Vesicatoria* (*Xcv*), the causal agent of bacterial spot disease (Moss et al., 2007), and the obligate biotrophic fungus *Oidium neolycopersici (On)*, the causal agent of powdery mildew in tomato (Jacob et al., 2008), upon silencing of *TOR*. We compared disease symptoms in L5 of *TOR* silenced and non-silenced plants after inoculated with *Xcv* or *On*. Leaves of *TOR* silenced M82 plants showed lower *Xcv* disease symptoms and exhibited a significant decrease in *On* disease severity in comparison to non-silenced plants. With the exception of *CKX*, none of the genotypes displayed a significant reduction in *Xcv* disease symptoms upon silencing (**Figure S4A**). In the case of *On* disease symptoms, *CKX* and *ga20ox* both showed a reduction in disease symptoms (**Figure S4B**).

### TOR mediates CK-driven developmental cues

Transgenic tomato lines with altered leaf CK content have altered developmental programs, resulting in quantifiable phenotypic changes in leaf development. *pBLS>>IPT7* has significantly more complex leaves, while *pFIL>>CKX*3 has significantly simpler leaves, when compared with their M82 background (Shani et al., 2010). Likewise, *pFIL>>PROΔ17* and *clausa* have more complex leaves (Israeli et al., 2021), while *pro^ΔGRAS^* has simpler leaves in comparison to the M82 background (Livne et al., 2015). To investigate whether TOR plays a role in GA and CK-mediated leaf development, we compared the leaf complexity of lines with different GA and CK levels upon *TOR* silencing. Interestingly, while the leaves of the WT M82 plants did not show any significant developmental changes in response to *TOR* silencing, we observed a reduction in leaf complexity in the highly complex *IPT, clausa*, *ga20ox3*, and *PROΔ17* plants, and an increase in leaf complexity in the simple-leaved *CKX* and *pro* plants **(Figure 3A-B)**, suggesting that TOR is required to execute the developmental cues generated by CK and GA. In addition, *TOR* silencing significantly promoted shoot length (plant height) in *IPT, clau, ga20ox3* and *PROΔ17*, while it reduced the height of *pro*. No significant difference in plant height was observed in *CKX*.

**Figure 3:**
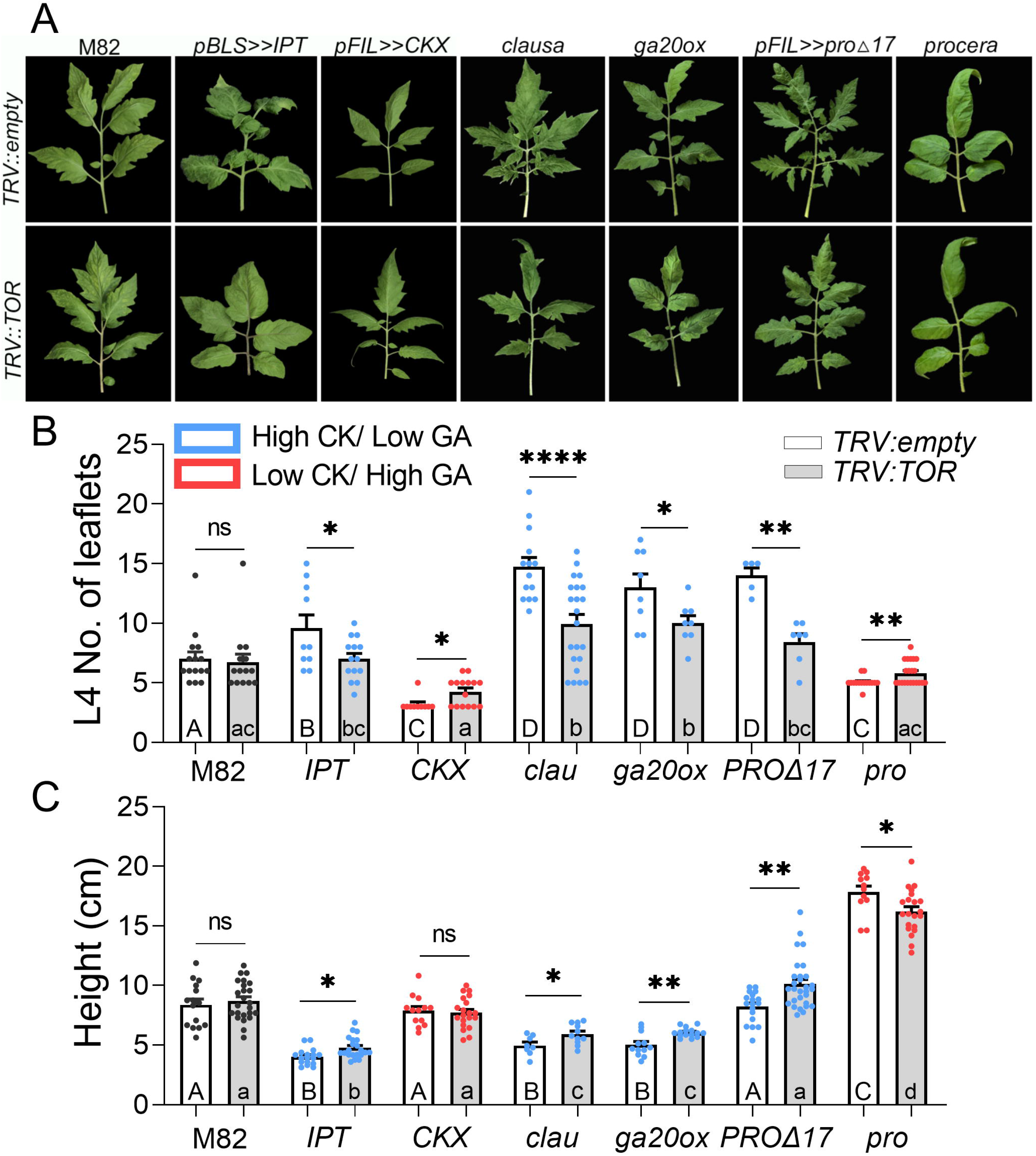
The CK/GA balance governs effects of *TOR* silencing on leaf development. *S. lycopersicum* plants of altered CK/GA genotypes: increased CK content *pBLS>>IPT7* (“*IPT*”), decreased CK content *pFIL>>CKX3* (“*CKX*”), increased CK sensitivity and decreased GA sensitivity *clausa* mutant (“*clau*”), decreased GA content mutant (“*ga20ox*”), decreased GA signaling *pFIL>>proΔ17* (“*PROΔ17*”), increased GA signaling *procera* (“*pro*”) and their WT background M82, were *TOR*-silenced using VIGS. 4 weeks after silencing, leaf complexity was quantified by counting the leaflets on leaves 4 (**A, B**), and height was measured (**C**). Experiment was conducted 3 times. Bars represent Mean ±SEM, all points shown. Asterisks indicate statistically significant changes in leaf complexity (B) or plant height (C) upon *TOR* silencing, and different letters indicate statistically significant differences among samples, upper case for control-silenced and lower-case for TOR-silenced, in Welch’s ANOVA with Dunnett’s post hoc test, or in student’s t-test with Welch’s correction. **B:** N>5 individual plants, **C:** N>8 individual plants. *p<0.05, **p<0.01, ***p<0.001, ****p<0.0001, ns-non significant.

To further examine the importance of TOR in mediating hormonal signals leading to abnormal tomato leaf phenotypes, we downregulated *TOR* in several classical tomato mutants, and examined the effect on leaf phenotypes. A dominant mutation in the TCP transcription factor *LA* (*LANCEOLATE*), known as *La2*, results in highly simple leaves, and overexpression of the *miR* that regulates *LA* expression, *miR*319 (as in *pBLS>>JAW*) causes highly complex leaves levels (Ori et al., 2007). *LA* was demonstrated to be involved in the regulation of the balance between CK and GA during leaf development (Israeli et al., 2021), and its activity is mediated in part by positive regulation of GA (Yanai et al., 2011) and negative regulation of CK (Efroni et al., 2013; Israeli et al., 2021). *CLAUSA* (**Figure 3**) and *LA* jointly regulate leaf development through the CK-GA balance (Israeli et al., 2021). The classical mutants *BIPPINATE* (*bip*) and *DOUBLE-DISSECTED LEAF* (*ddl*) were both previously found to be related to the KNOX-BELL machinery (Kimura et al., 2008; Nakayama et al., 2021). BELL proteins negatively regulate KNOX genes, and as such, can affect CK and GA levels (Bolduc & Hake, 2009; Jasinski et al., 2005; Sakamoto et al., 2001; Yanai et al., 2005). *POTATO-LEAF* (*c*) is a MYB transcription factor mutant shown to be involved in branching and boundary formation (Busch et al., 2011). *TOR* inhibition “normalized” the phenotypes observed in all these mutants (**Figure S5**), demonstrating TORs involvement in the mediation of hormonal signals and execution of development, in accordance with previous studies (reviewed in: McCready et al., 2020).

### GA-response is mediated by TOR

Exogenous GA can affect developmental programs and phenotypes (Jasinski et al., 2008; Sun, 2010). Given our results that TOR mediates hormonal signals, we hypothesized that *TOR* silencing would reduce GA sensitivity. Therefore, we next characterized the effect of treatments with different concentrations of GA on growth and immunity phenotypes of *TOR* silenced plants. GA-treated plants showed a concentration-dependent increase in *B. cinerea* disease susceptibility (**Figure 4A**) and plant height (**Figure 4B**), and a reduction in leaf complexity (**Figure 4C-D**) in comparison with untreated plants, as previously described (Fleishon et al., 2011). In response to GA treatment, *TOR* silenced plants showed a milder increase in disease susceptibility and plant height, and a milder reduction in leaf complexity, as compared with non silenced plants (**Figure 4A-D**). *TOR* silenced plants were more resistant to *B. cinerea* infection in the mock treatment, as previously described (Marash et al., 2022). Apart from plant height, which remained significantly differential among control and TOR-silenced plants, GA treatment at the highest concentration of 100 µM had similar effects on *TOR* silenced and non silenced plants (**Figure 4**). The reduced GA-sensitivity upon *TOR* silencing suggests that GA-response is mediated, at least in part, by TOR.

**Figure 4:**
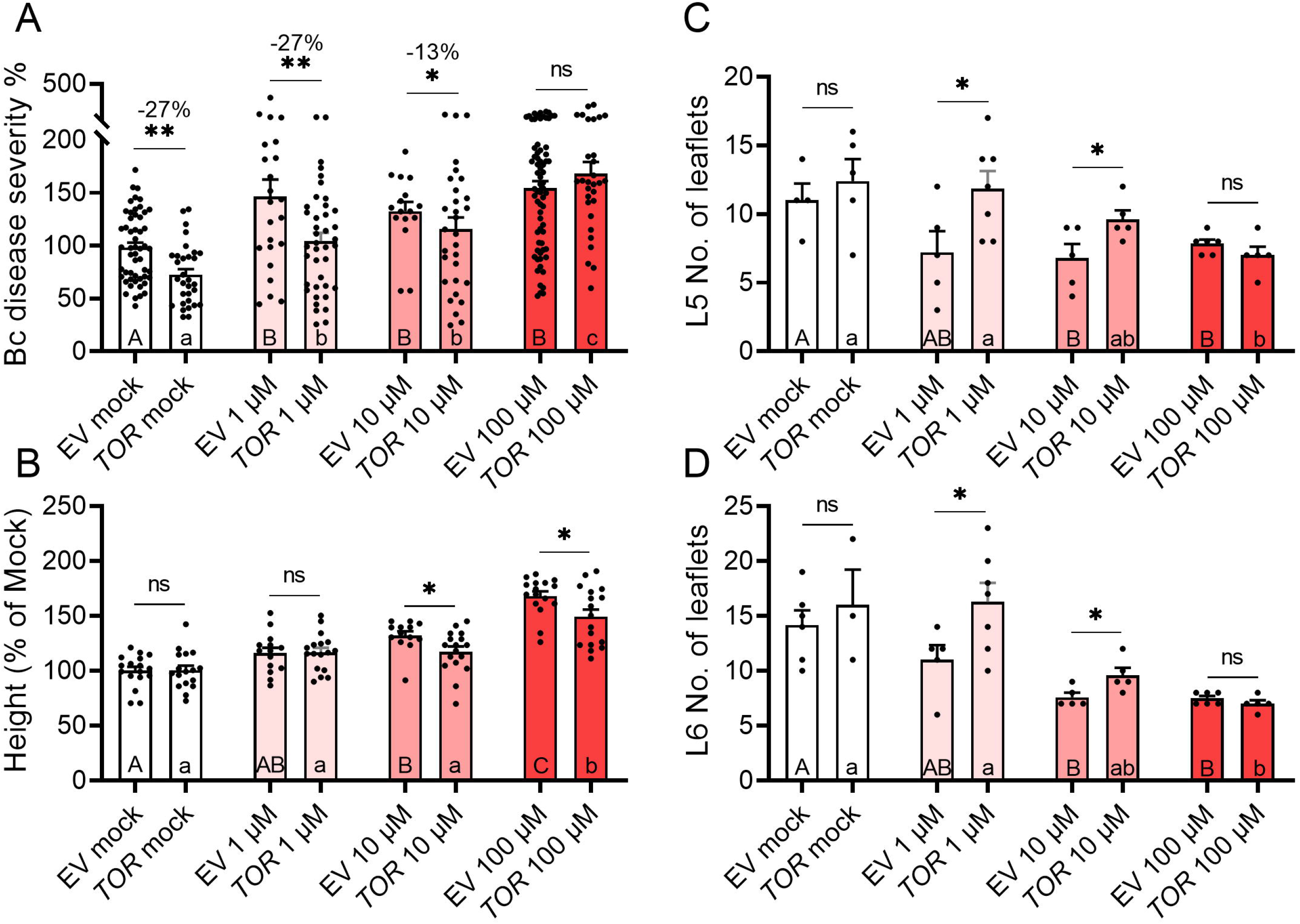
*TOR* silencing affects GA response. *S. lycopersicum* cv M82 plants were *TOR*-silenced using VIGS. One week after silencing, mock and *TOR*-silenced plants were treated with indicated concentration of GA3, three times a week, for 2 weeks, by spraying. 4 weeks after silencing, plants were challenged with *B. cinerea (Bc)* mycelia from a 72h old-culture (**A**), height was measured (**B**), and leaf complexity was quantified by counting the leaflets on leaves 5 (**C**) and 6 (**D**). Experiment was conducted 3 times. Bars represent Mean ±SEM, all points shown. Asterisks indicate statistically significant changes in B. cinerea disease (**A**), plant height (**B**), or leaf complexity (**C-D**) upon *TOR* silencing, and different letters indicate statistically significant differences among samples, upper case for control-silenced and lower-case for TOR-silenced, in Welch’s ANOVA with Dunnett’s post hoc test (**A**), one-way ANOVA with Bonferroni’s post hoc test (**B**), or in student’s t-tests with Welch’s correction (**C-D**). **A:** N=30, **B:** N=17, **C:** N=5, **D:** N>3. *p<0.05, **p<0.01, ns-non significant.

### TOR inhibition mediated immunity and CK mediated immunity do not augment each other

To further examine the relationship between TOR inhibition and CK in plant defense, we assessed *Bc* disease sensitivity of M82 plants upon Torin2 and 6-BAP treatment. Plants were treated either with Torin2 6-BAP, or both. As we previously reported (Gupta et al., 2020; Marash et al., 2022), both 6-BAP and Torin2 treatments promoted disease resistance, as lesion size was reduced by about 40% with either Torin2 or CK **(Figure S6A)**. Treatment with both Torin2 and 6-BAP, however, had no additive effect on disease resistance. To assess the effect of 6-BAP and Torin2 on plant defense responses, we analyzed ROS accumulation and ion leakage with or without Torin2 and 6-BAP treatment **(Figure S6B-C).** ROS accumulation and ion leakage were increased by 6-BAP and Torin2 treatments. However, no additive effect on induction of defense responses was observed upon combined treatment with both Torin2 and 6-BAP, suggesting that TOR inhibition and CK likely promote disease resistance through overlapping pathways.

### TOR inhibition alters CK and GA pathway genes and reduces CK response

Since *TOR* inhibition reduced the sensitivity to signaling cues mediated by CK and GA, we analyzed the effect of Torin2 on the expression of CK and GA metabolic and signaling genes. CK is synthesized by IPT enzymes (Kakimoto, 2001; Sakamoto et al., 2006; Takei et al., 2001), activated by LOG enzymes (Kurakawa et al., 2007; Kuroha et al., 2009), and perceived by a response regulator array (Argyros et al., 2008; Ishida et al., 2008). Type-A response regulators (*TRRs* in tomato) are known to be CK-responsive (Fleishon et al., 2011). The deactivation of CKs can happen either through conjugation or irreversible degradation by *Cytokinin oxidase/dehydrogenase* (*CKX*) (Mok & Mok, 2001; Werner et al., 2006). The expression of the CK-responsive response regulator genes, *TRR3/4* and *TRR16B* was downregulated by Torin2 treatment in developing leaves and unaffected in mature leaves, while the expression of the CK inactivating genes *CKX2* and *CKX5* was upregulated in developing leaves and downregulated in mature leaves (**Fig. 5A-B**). Expression levels of *TRR5/6/7* and *CKX6* was unaffected in both tissues. Expression of the CK-biosynthesis genes *IPT5* and *IPT6* was downregulated in developing leaves upon Torin treatment, and unaffected in mature leaves. *IPT3* was unaffected in developing leaves and downregulated in mature leaves (**Figure 5C-D**). Expression of the CK activating LOG enzymes *LOG5* and *LOG8* was reduced by Torin in developing leaves (**Figure 5C-D**). Torin also induced *LOG8* expression in mature leaves. *LOG4* was unaffected in both tissues. Overall, TOR inhibition causes CK biosynthesis, CK activation, and CK signaling, to be inhibited in developing leaves, and somewhat promoted (apart from *IPT3*) in mature leaves.

**Figure 5:**
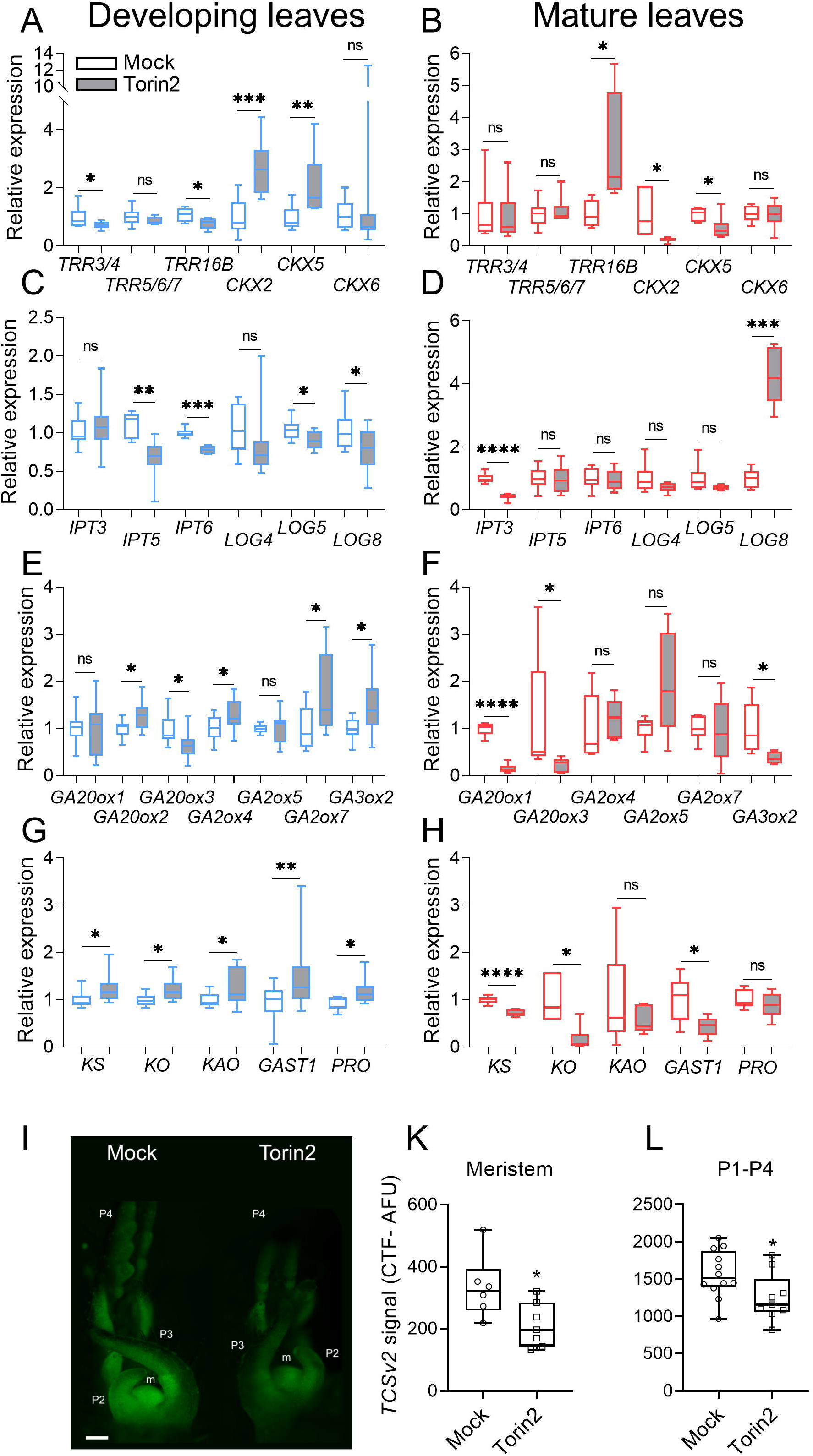
*TOR* inhibition alters CK and GA pathway gene expression and reduces CK response in the meristem of young shoots. **A-H**: Gene expression analysis of the indicated CK (A-D) and GA (E-H) pathway genes, with and without Torin2 (2 µM) treatment, was measured by RT-qPCR. 1:5000 DMSO in DDW served as Mock. Relative expression was calculated using the geometric mean of the gene copy number obtained for three reference genes, and normalized to the expression following Mock treatment. Developing leaves consisted of 3-week-old shoot apexes with 6 youngest primordia, mature leaves the fifth leaf of 6-week-old plants. The following normalizer genes were used: for developing leaves: *RPL8* (Solyc10g006580), *EXP* (Solyc07g025390), and *CYP* (Solyc01g111170), and for mature leaves, *RPL8*, *CYP*, and *Actin* (Solyc11g005330). Analysis was conducted on 6 individual plants. Boxplots represent inner quartile ranges (box), outer quartile ranges (whiskers), median (line in box). Asterisks indicate significant differential regulation upon Torin2 treatment in Welch’s t-test comparing each gene, *p<0.05, **p<0.01, ***p<0.001, ns-non significant. A-B: Tomato response regulators (TRRs) and CK oxidases (CKXs). C-D: Iso-pentenyl transferases (IPTs) and lonely-guy CK activating enzymes (LOGs). E-F: GA20, GA2, and GA3 oxidases. G-H: GA biosyntesis upstream enzymes (KA, KO, KAO) and responsive genes (GAST1, PRO). **I-L**: *S. lycopersicum* cv. M82 10 day-old seedlings expressing VENUS driven by the cytokinin responsive promoter TCSv2 were treated with Torin2 (2 µM) or Mock (1:5000 DMSO in DDW) for 48 h. **I** Typical Mock treated and Torin2 treated shoots are depicted. Images captured under identical conditions. The meristem (m), second (P2) third (P3) and fourth (P4) youngest leaf primordia are indicated. Bar-1000 µM. TCSv2 driven total Venus fluorescence in the meristem (**I-K**) or leaf primordia (P1-P4) (**I-L**) was measured as corrected total fluorescence (CTF), in images captured under identical conditions. Boxplots represent inner quartile ranges (box), outer quartile ranges (whiskers), median (line in box), all points shown. Asterisks indicate significant TCSv2 signal reduction upon Torin2 treatment in an unpaired two-tailed *t*-test, N>7, *p<0.05.

The production of GA from Geranylgeranyl diphosphate (GGDP) requires multiple enzymes, including *ent*-kaurene synthase (*KS*), *ent*-kaurene oxidase (*KO*), *ent*-kaurenoic acid oxidase (*KAO*), GA20 oxidase (*GA20ox*), and GA3 oxidase (*GA3ox*). The GA metabolic genes *GA20ox* and *GA3ox* convert precursors of GA into its active form through a series of oxidation reaction, while the *GA2ox* genes are essential for inactivating GA (Hedden, 2020). We found that the expression of key genes involved in the biosynthesis of active forms of GAs, *GA20ox2 and GA3ox2*, was upregulated by Torin2 in developing leaves (**Figure 5E**). *GA3ox2* was downregulated by Torin2 in mature leaves, and *GA20ox1* was unaffected in developing leaves and downregulated in mature leaves, while *GA20ox3* was downregulated in both developing and mature leaves (**Figure 5E-F**). The expression of the GA-inactivating enzymes *GA2ox4* and *GA2ox7* was upregulated in developing leaves and unaffected in mature leaves. (**Figure 5E-F**). The expression levels of genes involved in early GA biosynthesis, *KS*, *KO*, and *KAO,* was elevated by Torin2 in developing leaves, and reduced in mature leaves (except for *KAO* that was unaffected in mature leaves) (**Figure 5G-H**). Likewise, the expression of the GA response and signal transduction genes *PROCERA* and *GAST1* (Shi & Olszewski, 1998) was upregulated in developing leaves, while *GAST1* was downregulated in mature leaves **(Figure 5G-H)**. Overall, TOR inhibition causes GA biosynthesis, GA activation, and GA signaling, to be promoted in developing leaves, and inhibited in mature leaves. These results suggest the existence of a feedback regulatory mechanism between GA, CK and TOR that may help plants to fine-tune the level of these hormones upon pathogen attack.

To further examine the effect of TOR inhibition on CK signaling in young developing shoots, we used the CK-response transgenic reporter line *pTCSv2::3xVENUS*, that expresses *VENUS* under the control of the CK-responsive synthetic promoter TCSv2 (Steiner et al., 2016, 2020; Zürcher et al., 2013). We found that in the presence of Torin2, there is a reduction in CK signaling. Torin treated *pTCSv2::3xVENUS* shoots showed a significant reduction in VENUS signal relative to untreated shoots, in the meristem (**Figure 5I, K**) and four youngest leaf primordia (**Figure 5 I, L**). Similar results were achieved in TCS expressing plants in which *SlTOR* was silenced by VIGS **(Figure S7)**. This aligns with the reduction in *TRR3/4*, *TRR16B*, *IPT5*, and *IPT6*, and the increase in *CKX*2 and *CKX5* expression in developing leaves upon Torin2 treatment (**Figure 5A, C**).

Pathogen infection has been previously demonstrated to alter the expression of CK pathway genes (Argueso et al., 2012; Gupta et al., 2020; Gupta, Leibman-Markus, et al., 2021). To investigate possible effects of TOR status on pathogen-driven alterations in CK pathway genes, we examined gene expression in L5 of 10-week-old WT M82 plants 24h following spray-inoculation with *Bc* spores, with and without Torin2 treatment by petiole feeding (**Figure S8**). CK-mediated signaling (examined by assaying expression of the response regulators *TRR3/4* and *TRR5/6/7*) showed the expected increase following Bc treatment (**Figure S8A-B**). *TRR3/4* was reduced by Torin2 treatment, and following Bc and Torin2 co-treatment, returned to background levels (**Figure S8A**). *TRR5/6/7* was not affected by Torin2, however, upon combined treatment, it increased beyond the levels elicited by *Bc* alone (**Figure S8B**), suggesting that TOR mediated effects on the CK pathway may be differential through interactions with different CK pathway genes. *CKX2* and *CKX5* expression was increased in response to Torin2 or *Bc* infection, and the combination of both *Bc* infection and Torin2 treatment did not further augment this increase in their expression (**Figure S8C-D**).

### CK and GA can affect TOR activity

To further investigate a possible feedback regulatory mechanism between GA, CK and TOR, we examined how CK and GA might modulate TOR activity. To do so, we analyzed the phosphorylation status of S6 kinase 1 (S6K1), a conserved TOR substrate that has been previously used as an indication of TOR activity in plants (Cao et al., 2019; Li et al., 2017; Liu et al., 2021; Song et al., 2022; Xiong & Sheen, 2012; Ye et al., 2022). We treated plants with 100 µM GA and CK, separately or simultaneously, and tested the phosphorylation status of S6K1 after 4 hours, in both developing and mature leaves. As shown in **Figure 6**, exogenous CK treatment reduced the phosphorylation of S6K1 by TOR in both developing and mature leaves, suggesting that CK may negatively regulate TOR activity. By contrast, GA treatment, as well as the GA and CK co-treatment, significantly increased the level of TOR-phosphorylated S6K in developing and mature leaves in comparison to mock, suggesting that GA can inhibit CK-mediated reduction of TOR activity.

**Figure 6:**
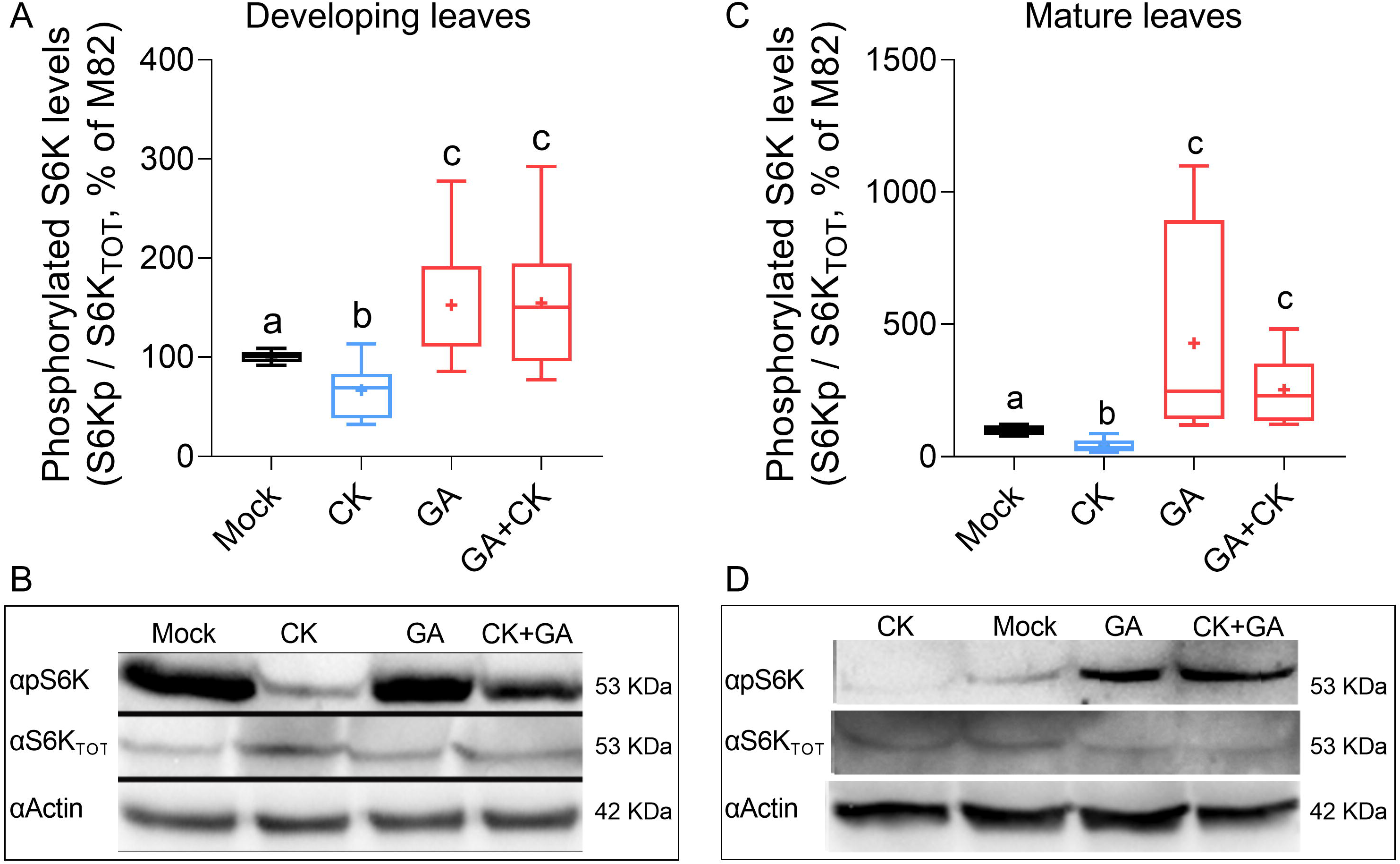
CK and GA affect TOR activation. *S. lycopersicum* cv. M82 6-week-old plants were treated with Mock (10 µM NaOH), 10 µM of the CK 6-benzylaminopurine (6-BAP), or 10 µM of GA3. 24 h after treatment, total cellular proteins were prepared from developpoing (**A-B**) and mature (**C-D**) leaves. Developing leaves consisted of 3-week-old shoot apexes with 6 youngest primordia (10-12 plants per biological repeat), and the fifth leaf of 6 individual plants 6-week-old plants was used for mature leaves. TOR activation was expressed as the ratio between phosphorylated S6K and total S6K, detected using specific antibodies. Actin was detected as an additional control. Experiment was repeated two independent times, A-B: N=8, C-D: N=6. Boxplots represent inner quartile ranges (box), outer quartile ranges (whiskers), median (line in box), mean (“+” sign). Different letters indicate statistically significant differences among samples in a Mann Whitney U test, A: p<0.045, C: p<0.0019.

We found that leaf developmental status affected disease resistance (**Figure 1**) and TOR inhibition-mediated gene expression (**Figure 5**). Therefore, we tested whether TOR-inhibition mediated immunity could be dependent on leaf developmental-stage. Interestingly, *Bc* susceptibility decreased with leaf age (**Figure 1A, 7A**). While Torin2 reduced *Bc*-induced disease in L3 and L5, disease level of L8 treated with Torin2 was similar to that observed in untreated leaves (**Figure 7B**). This finding demonstrates that the extent of TOR-mediated immunity increases with leaf developmental age.

**Figure 7:**
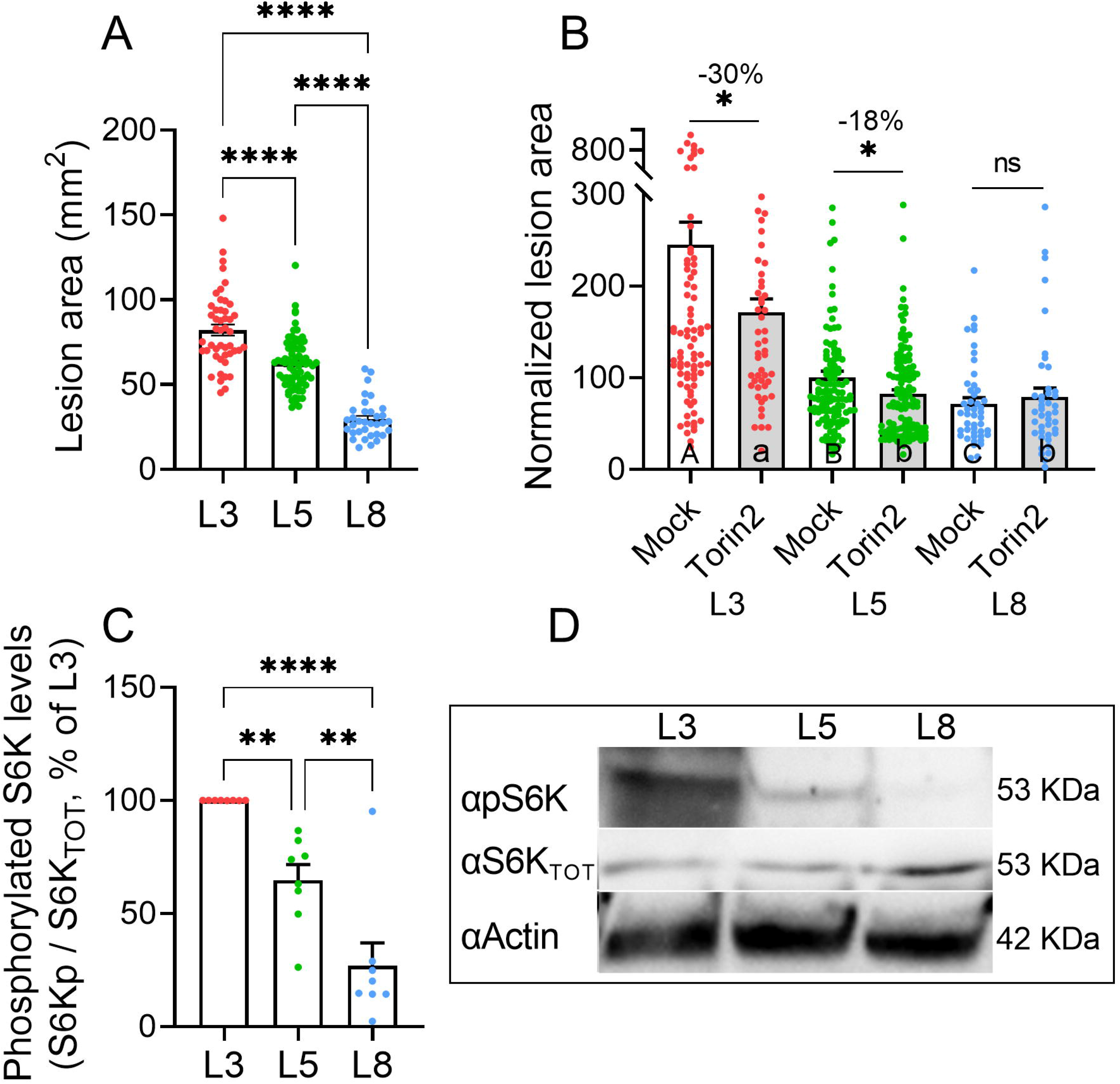
*TOR* activity and *TOR* inhibition mediated disease resistance depend on leaf developmetal stage. **A**: Different leaves as indicated from *S. lycopersicum* cv. M82 5 week old plants were infected with *B. cinerea*. Asterisks denote staistical significance among indicated samples in Welch’s ANOVA with Dunnett’s post hoc test, N>35, ****p<0.0001. Box plots represent inner quartile ranges (box), median (line in box), and outer quartile ranges (whiskers), all points shown. **B:** Different leaves as indicated from *S. lycopersicum* cv. M82 5 week old plants, were treated with Mock (1:5000 DMSO in DDW), or 2 µM Torin2. Plants were challenged with *B. cinerea (Bc)* mycelia from a 72h old-culture 24 h after treatment. Bars represent mean ±SEM, all points shown. Experiments were repeated 3 independent times. Asterisks indicate statistically significant decreases in B. cinerea infection upon Torin2 treatment as compared with Mock treatment, and letters indicate statistically significant differences among samples, upper case letters for Mock treated genotypes and lower case letters for samples treated with Torin2, in Welch’s ANOVA with Dunnett’s post-hoc test, N>40, *p<0.05. Percentage of disease reduction is indicated above asterisks. **C-D**: Total cellular proteins were preapared from the indicated leaves of *S. lycopersicum* cv. M82 5 week old plants. TOR activation was expressed as the ratio between phosphorylated S6K and total S6K, detected using specific antibodies. Actin was detected as an additional control. Experiment was repeated 4 times, N=8. Boxplots represent inner quartile ranges (box), outer quartile ranges (whiskers), median (line in box), mean (“+” sign), all points shown. Asterisks indicate statistically significant differences among indicated samples in One-way ANOVA with Tukey’s post hoc test, **p<0.01, ****p<0.0001. D: Representative blots.

Given the results that both CK and GA affect TOR activity (**Figure 6**), and the fact that the ratio between CK and GA changes during leaf development (Weiss & Ori, 2007), we examined how TOR-kinase activity changes throughout leaf maturation. TOR activity was lowest in L8, the youngest, most developmentally morphogenetic leaf, and gradually increased with age, with the highest activity observed in the mature, no longer morphogenetic L3 (**Figure 7C-D**). This is in agreement with earlier report demonstrating that TOR’s activity level increases with leaf age in Arabidopsis (Brunkard et al., 2020).

## Discussion

The TOR pathway senses many different inputs such as changes in cellular energy status, hormone levels, light, and abiotic or biotic stresses, to regulate growth, metabolism, transcription, and translation (Dobrenel et al., 2016). Various reports have suggested that TOR balances between growth and defense responses in plants (Caldana et al., 2019; De Vleesschauwer et al., 2018; Margalha et al., 2019; Ryabova et al., 2019). Previous work, for example, indicated that in plants, TOR acts as a molecular ‘switch’ at the intersection of growth and defense, and activates cell proliferation and plant growth at the expense of defense (De Vleesschauwer et al., 2018). Consistent with this, we have recently reported that plant immunity and defense responses are enhanced upon TOR downregulation in tomato (Marash et al., 2022).

### TOR mediates plant immunity

Despite the extensive research on the role of the GA signaling in plant growth and development, there has been limited study on its role in plant defense responses (Bari & Jones, 2009; Wang et al., 2013). Previous works in Arabidopsis show that treatment with GA increases the resistance to (hemi)biotrophic bacterial pathogens, but reduces resistance to necrotrophic pathogens (Navarro et al., 2008), whereas in rice, it increases resistance to necrotrophic pathogens, and reduces resistance to (hemi)biotrophic pathogens (de Vleesschauwer et al., 2012, 2016; Qin et al., 2013; Yang et al., 2008). This implies that the effect of GA on plant immunity depends on both the host plant and type of pathogen involved (reviewed in: De Bruyne et al., 2014). As shown here (**Figure 4**), GA likely affects plant immunity in a similar manner in tomato and *Arabidopsis*. Our data suggest that immunity mediated by CK or GA requires inhibition of TOR activity for execution. This could also explain why TOR inhibition did not further enhance defense responses or *Bc* resistance when combined with CK application **(Figure S6)**. In tomato, TOR inhibition (Marash et al., 2022) or high endogenous CK levels (Gupta et al., 2020), promote pathogen resistance in a SA-dependent manner. Thus, a possible mechanism that could explain our results is that TOR and CK signaling pathways coordinately regulate plant defense responses through the modulation of SA. We hypothesize that the crosstalk between TOR and CK signaling could be involved in the ability of plants to modify growth and developmental programs upon pathogen attack, and thus enable a faster activation of plant defense responses. Conversely, the result that TOR downregulation decreased *Bc*-sensitivity in high GA (or low CK) genotypes or upon exogenous GA treatment, whereas it had no significant effect in low GA (or high CK) genotypes, suggests that TOR and the CK and/or GA pathways might share signaling components.

### TOR mediates the leaf developmental program

Leaf development relies on the balance between GA, which promotes differentiation, and CK, which promotes morphogenesis (Hay et al., 2005; Jasinski et al., 2005; Yanai et al., 2005). Thus, CK and GA have a partial antagonistic role in leaf development (Bar et al., 2016; Fleishon et al., 2011). Generally, GAs are considered as differentiation promoting hormones which help to complete developmental programs and regulate the achievement of final organ forms. GA shortens the morphogenetic stage of leaf development by promoting differentiation (Shwartz et al., 2016). The termination of the juvenile phase is associated with an increase in the levels of endogenous GA. This suggests that GAs promote the transition from a juvenile- or developing-state, to an adult- or differentiated-state (Porri et al., 2014). On the other hand, CKs are known to alter leaf development and morphology (Hay & Tsiantis, 2010; Shani et al., 2010; Werner et al., 2003), and are regarded as factors that promote “juvenility” by promoting morphogenesis and delaying differentiation and senescence (Shwartz et al., 2016). Dividing tissues in the leaf have the highest levels of cytokinin, while bioactive gibberellins peak at transition zone between the division and expansion zone (Nelissen et al., 2012).

It has been suggested that active growth and cell-cycle progression are required for the formation of the hormonal axis, involving auxin and cytokinin, that is required for organ formation and patterning during plant development (Du et al., 2018). Accordingly, several studies reported that TOR plays a role during leaf development in *Arabidopsis*. For example, *TOR* downregulation has been shown to result in the production of smaller leaves with fewer cells (Caldana et al., 2013) whereas *TOR* overexpression results in the production of bigger leaves with larger cells (Deprost et al., 2007). Likewise, mutation in *AtLST8*, a member of the TOR complex, results in a reduction in the number of leaves and in leaf size (Moreau et al., 2012), and mutation in *AtRAPTOR1B*, another component of the TORC1 complex, stalls leaf initiation (Anderson et al., 2005). By contrast, we did not observe any significant phenotypic alterations in the tomato WT M82 cultivar upon *TOR* silencing. This could be ascribed to the different inhibition methods used, or the different plant species. While TOR inhibition might affect the translation of the ectopically expressed proteins in transgenic lines, given similar results achieved with several mutants, this would be unlikely to explain our results, however, it should also be noted that our developmental analyses could be limited by use of the VIGS system. Improved systems to comprehensively study plant development upon TOR inhibition will no doubt emerge in the future.

Our work indicates that TOR is required for the developmental response to hormonal signals. Response to exogenous GA treatment, as well as the patterning of leaf organs programmed by CK and GA, were perturbed by TOR inhibition. These findings agree with a previous report demonstrating that mutants in *raptor*, a protein in the TOR complex, were less sensitive to exogenous GA treatment (Zhang et al., 2018). The increased leaf complexity in lines with a high CK/GA ratio, and decreased complexity in lines with a low CK/GA ratio, were both partially rescued to WT M82 levels as a result of *TOR* silencing (**Figure 3**). In general, TOR silencing partially rescued a variety of aberrant leaf developmental phenotypes (**Figures 3, S5**). This brings forth the notion that TOR mediates signals from additional hormones, or that TOR is responsible for the reduction to practice of a variety of cues and signals generated by the balance and crosstalk of several developmental hormones (Greenboim-Wainberg et al., 2005; Israeli et al., 2021). The signaling events that are transmitted from the CK/GA ratio and encode leaf complexity, are no longer properly transmitted or decoded when TOR is attenuated. Thus, TOR supports hormonal signal output, resulting in the typically observed leaf phenotypes. When TOR is inhibited, the signaling output from the aberrant hormonal balance in developmental mutants is no longer supported by TOR, resulting in milder phenotypes. These phenotypic changes could indicate that CK and GA distribution and/or signaling are altered in response to *TOR* silencing, or that factors which execute organ patterning downstream of hormonal cues are dependent on TOR status. It is therefore possible that TOR is required for the execution of a variety of cues that are integrated to form a cohesive leaf developmental program. Notably, TOR inhibition reduced CK signaling and increased GA signaling in developing leaves, and promoted CK signaling and reduced GA signaling in mature leaves (**Figure 5**). Due to the lethality of complete TOR inhibition, and the limitations of the VIGS system, we were not able to assess a full leaf developmental time course upon TOR inhibition. However, we would expect TOR inhibition early in leaf development to result in precocious differentiation and simpler leaves, while TOR inhibition in late development could potentially lengthen the morphogenetic window.

The TOR pathway has been implicated in the regulation of translation and ribosome biogenesis in mammals and plants, and therefore its activity is tightly regulated (Pereyra et al., 2020). Interestingly, proteome analysis of CK activity in *Arabidopsis* demonstrated extensive differential regulation of ribosomal proteins in response to CK (Brenner & Schmülling, 2012). In another proteomic study, the functional classification ‘Ribosome biogenesis’ was found to be strongly differential in response to CK depletion or overproduction (Černý et al., 2013). Thus, the molecular mechanisms underlying the effect of CKs on leaf development and morphology could potentially be mediated by changes in translational processes due to differential regulation of ribosomal proteins. Horiguchi et al., 2011 demonstrated that ribosomal proteins play a key role in *Arabidopsis* leaf development, supporting this notion. Thus, it is likely that TOR is involved in the execution of CK-mediated signals, in both defense and development, by its direct or indirect regulation of proteins required for the execution of these processes.

### Immunity to B. cinerea depends on developmental status

The phenomenon in which developing leaves show stronger resistance than developmentally mature leaves (**Figures 1, 7**) has been described as “leaf-stage associated disease resistance” (Berens et al., 2019; Develey-Rivière & Galiana, 2007; Xu et al., 2018). Zhang & Chen, 2009, for example, showed that as tomato leaves age, they become more susceptible to diseases caused by *B. cinerea* and *Fusarium oxysporum*. In *Arabidopsis*, it was proposed that this resistance is due to higher accumulation of SA in young leaves (Zeier, 2005). Our results suggest that high CK or low GA signals prevent the increase in *B. cinerea* susceptibility that normally occurs with developmental stage progression. As young organs have a higher CK/GA ratio than mature organs, and since the CK/GA balance, rather than the content or signal of each individual hormone was reported to execute leaf developmental functions (Fleishon et al., 2011; Shani et al., 2006; Shwartz et al., 2016), it is possible that leaf developmental stage-related *B. cinerea* susceptibility may also depend on the CK/GA balance. This idea corresponds to previous work demonstrating that altered GA levels were not able to prevent the age-related decrease in JA-mediated immunity (Mao et al., 2017). Therefore, disease resistance related to developmental status could potentially be attributed to the CK/GA balance, and further, be mediated by TOR in a manner similar to that we observed for leaf developmental processes. Thus, it emerges from our results that TOR activity supports GA-mediated processes and reduces disease resistance, while a decrease in TOR activity supports CK-mediated processes and increases disease resistance.

### Increased B. cinerea resistance in young leaves could be a result of decreased TOR activity

TOR’s low activity in young leaves (**Figure 7**) could account for their enhanced *Bc* resistance. The observation that CK application results in a decrease in TOR activation whereas GA application results in an increase in in TOR activation could potentially provide mechanistic insight for how resistance is affected by developmental-status. This indicates that resistance may be depended on TOR activity as well as the relationships between hormonal signals. Additionally, this implies the existence of a feedback mechanism that balances between TOR activity and these hormonal pathways. It is noteworthy that previous work in *Arabidopsis* cell suspensions revealed that kinetin and auxin both induced the phosphorylation of AtS6k (Turck et al., 2004), whereas the activation of *triticum aestivum* TOR (TaTOR) was induced during GA-triggered germination (Smailov et al., 2020). Our data demonstrate that the ability of CK to induce immunity is related to TOR status (**Figures 1, 7**), and that increased CK leads to a reduction in TOR activity (**Figure 6**). Our results align with those obtained in wheat, and differ from those obtained in *Arabidopsis*, possibly due to the use of cell suspensions, or due to differences among plant hosts.

#### The involvement of TOR in growth-defense tradeoffs

Recent studies demonstrating that growth and defense can be uncoupled suggest that resource reallocation toward immunity is not the sole factor of growth inhibition (Campos et al., 2016; Kliebenstein, 2016; Shchneider 2023). Phytohormones serve as clear mediators of the tradeoff between growth and defense, as they mediate development and defense responses (Berry & Argueso, 2022).

Leaves become more susceptible to necrotrophic diseases with age (**Figure 1**). This increased susceptibility appears to be largely influenced by the CK/GA ratio, which correlates with leaf developmental stage. Based on that observation, we speculate that this might be a mechanism by which old plants, in which the CK/GA ratio is very low, die and allow for the allocation of resources to their offspring. Thus, the change in CK/GA ratio in mature plants is translated into defense hormonal outputs that modulate plant immunity.

Following our findings, we propose a hypothetical model by which TOR modulates CK and GA signaling and acts as a mediator of both developmental and defense processes, potentially regulating development-defense trade-offs (**Figure 8**). Developing leaves have low TOR activity (Brunkard et al., 2020, **Figure 7**), a relatively high CK to GA ratio (Shwartz et al., 2016), and are resistant to *Bc*. Mature differentiated leaves have higher TOR activity, a lower CK to GA ratio, and are more sensitive to *Bc*. Our results suggest that developmental-status related resistance could depend on processes by which TOR transduces signals derived from the CK/GA balance. In mature leaves under standard growth conditions, TOR activity is high (the CK/GA ratio is relatively low), and morphogenesis is largely concluded. The “price” for this is relative disease susceptibility. When pathogens attack, CK increases and GA decreases (Meldau et al., 2012; Gupta et al 2020), and TOR becomes less active (Margalha et al., 2019), promoting disease resistance. The “price” for this disease resistance could be an arrest of growth, until the pathogen is vanquished. The relationship between TOR, GA and CK likely involves complex feedback mechanisms that are based on mutual regulation between these pathways, allowing for a gradual shift between growth and defense that should be addressed in future research.

**Figure 8:**
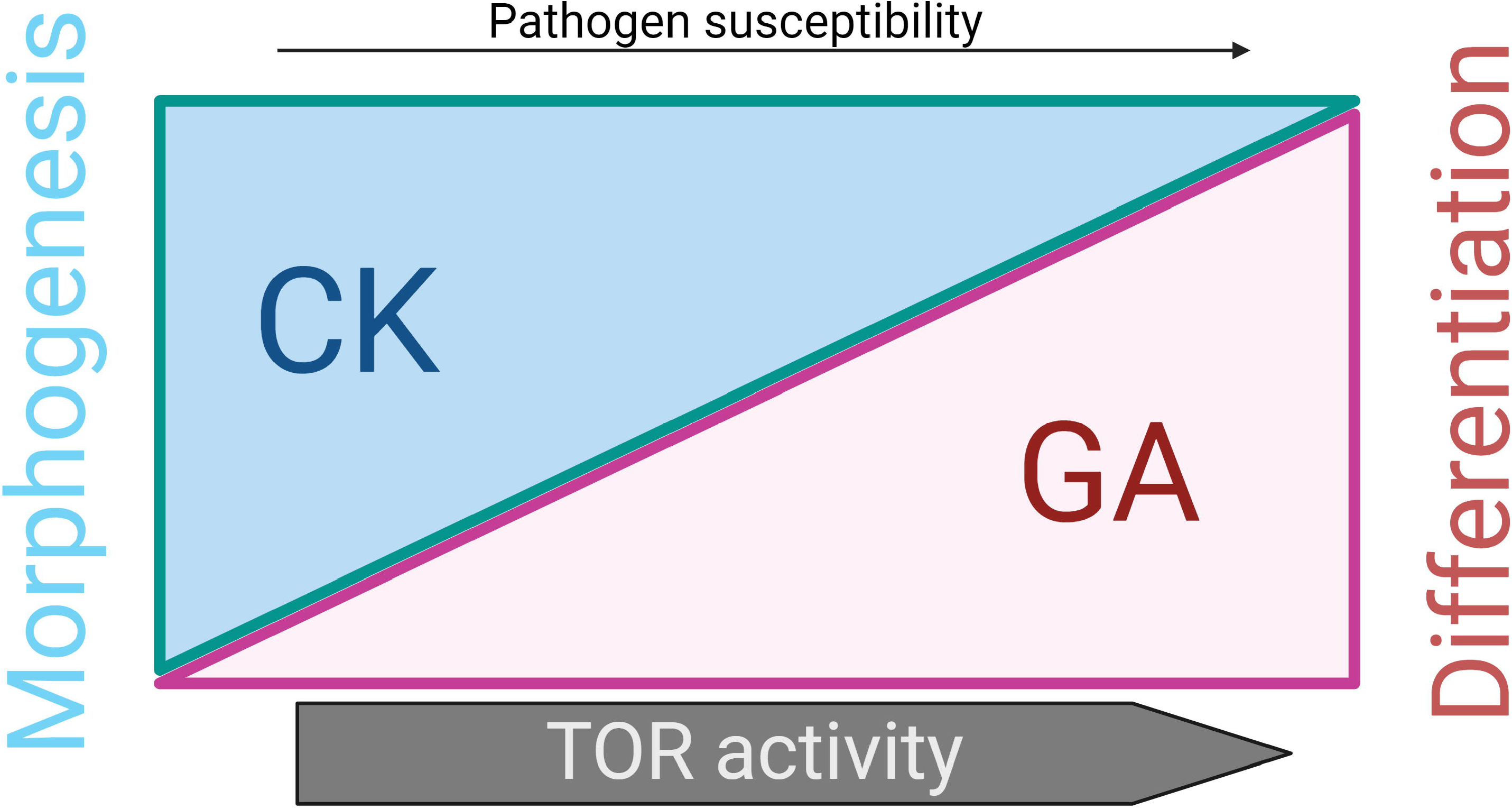
Model describing the interplay between the CK/GA balance and *TOR* in the regulation of development and defense cues. Cytokinin promotes both moprhogenesis and defense. Balanced CK/GA levels are required to achieve “normal” developmental patterning and disease resistance (**Fig.s 1-3**). In young leaves undergoing morphogenesis, CK signaling is high and TOR activity is low (**Fig.s 5, 7**), resulting in decreased pathogen susceptibility (Fig. 1). As leaves mature developmentally, CK signals are reduced and GA signals increase, the leaf morphogenetic potential declines as it differentiates, and TOR activity and pathogen suceptibility increase (**Fig.s 1, 7**). The CK/GA balance underlies both the morphogenetic potential and the disease susceptibility of the leaf. Inhibition of TOR results in rescue of altered CK/GA balances, partially restoring baseline developmental patterning and disease resistance (**Fig.s 3, 7**), and suggesting that TOR mediates these develpmental and defense cues originating from the CK/GA balance. In the tradeoff between development and defense, high CK can cause downregulation of TOR (Fig. 6), resulting in a shift towards defense (**Fig.s 1-2**), while high GA results in up-regulation of TOR activity (Fig. 6), and increased disese susceptibility (Fig. 4). The switch between development and defense may be modulated by cross-talk between environmental sensing and TOR status.

TORs involvement in both development and defense poises TOR as a prime regulator of tradeoffs between these two important aspects of plant life. Here, we demonstrate that CK and GA can regulate TOR activity, while TOR is required for the interpretation of defense and developmental signals originating from the CK/GA balance. The interaction between TOR, GA and CK could potentially help plants mediate growth and defense tradeoffs to adapt to the environment, with TOR sensing integrating environmental cues and stresses with plant hormonal balances, potentially allowing gradual shifts between growth and defense as a mechanism regulating plant robustness and survival under changing environments and pathogen loads. The relationship between TOR, GA and CK appears to involve complex feedback mechanisms based on mutual regulation between these pathways. It will be interesting to investigate how different metabolic states and biotic and abiotic stresses alter the cross talk between CK, GA and TOR in the future.

## Supporting information

All Supplemental Figures and Table in one file

## Acknowledgements

We thank members of the Spiegelman and Teper groups for technical assistance, and members of the Bar group for continuous support.

## Author Contributions

MB and IM conceived and designed the study. IM, ML-M, RG, GA, NL, DN, AI and MB formulated the methodology and carried out the experiments. IM, ML-M, RG, GA, and MB analyzed the data. All authors contributed to the writing of the manuscript.

## Competing Interests Statement

The authors declare no competing interests.

## Materials and methods

### Plant material and growth conditions

Plants were grown in soil (Green 332; Even-Ari Green, Ashdod, Israel) in a growth chamber set to long-day conditions (16/8 light/dark) at 24°C, or in a greenhouse under natural day length conditions.

The genotypes used in this study are detailed in the following table. Promoter line selection and hormone content are explained below.

#### Lines with altered GA signaling

the N’-terminal region of DELLA proteins contains the DELLA domain, which is required for the interaction with the GID1 receptor. Gain-of-function (GOF), dominant mutations in the DELLA domain block the interaction between DELLA and GID1 and prevent DELLA degradation. (Locascio et al., 2013; Murase et al., 2008). The C’-terminal region of DELLA interacts and represses multiple growth-promoting transcription factors (Locascio et al., 2013; Sun et al., 2012). Loss-of-function (LOF), recessive mutations in DELLA’s C’-terminus are linked to constitutive GA responses (Achard et al., 2006, 2008; Nir et al., 2017). DELLA proteins promote Jasmonic Acid (JA) signaling and repress salicylic acid (SA) biosynthesis and signaling, promoting susceptibility to biotrophs and resistance to necrotrophs. A DELLA loss-of-function mutant (quadruple *della* mutant, lacking four out of the five *Arabidopsis* DELLA proteins) for example, was shown to be partially insensitive to gene induction by JA, whereas a DELLA gain-of function mutant *gai* (constitutively active dominant DELLA mutant) was shown to be more sensitive (Navarro et al., 2008). There is only one DELLA protein in tomato (Livne et al., 2015). We used the GA deficient mutant *ga20ox3* and the DELLA gain-of-function line *pFIL>>GFP-PROΔ17* as lines with low GA signaling, and the DELLA loss-of-function mutant *procera^ΔGRAS^* as a line with high GA signaling. The gain-of-function mutant *pFIL>>GFP-PROΔ17* expresses the stable DELLA mutant protein *PROΔ17*, which lacks the DELLA domain. GA responses are constitutively suppressed in this mutant, resulting in a severe GA-deficient phenotype and GA insensitivity (Nir et al., 2017). *pro^ΔGRAS^* is a *procera* null mutant that lacks the entire C′-terminal region of DELLA and exhibits enhanced GA responses (Livne et al., 2015). The GA deficient mutant *ga20ox3* was generated using CRISPR/CAS9 essentially as described in Israeli et al., 2019, using the same constructs and methodology. gRNA primers are detailed in Table S1.

#### Lines with altered CK signaling

We used the high endogenous CK content genotype *pBLS >> IPT7*, which overexpresses the CK biosynthesis gene *ISOPENTYL TRANSFERASE 7 (IPT7)* and the low CK content genotype *pFIL>> CKX3*, which overexpresses the CK degrading enzyme *CKX OXIDASE3 (CKX3)* (Shani et al., 2010). We also used the increased CK sensitivity and decreased GA sensitivity mutant *clausa*. *CLAUSA* (*CLAU*) is a MYB transcription factor that promotes the transition from morphogenesis to differentiation by negatively affecting CK signaling and promoting GA signaling (Bar et al., 2016; Israeli et al., 2021).

### Rationale for genotype selection and reported CK and GA content in different genotypes

We used transgenic lines expressing genes of interest from the leaf specific promoters *FIL* and *BLS*. In tomato, the Arabidopsis *FIL* (filamentous flower) promoter drives expression throughout leaf primordia, starting from initiation (first plastochron), in initiating leaflets, and the abaxial side of the leaves. The *BLS* promoter drives expression later in leaf development, in primordia from about the fourth plastocrhon stage, and in young leaves (Lifschitz et al., 2006). Plants overexpressing *IPT* have been shown to contain increased levels of cytokinin many times, in various plant species (Márquez-López et al., 2019; Redig et al., 1996; Smigocki & Owens, 1988). Strongly increasing CK levels throughout the plant also led in some cases to undesirable phenotypes such as reduced apical dominance, increased lateralization, late flowering, and infertility. Therefore, tissue specific promoters were used to express IPT (Bartrina et al., 2011; Shani et al., 2010; Smigocki et al., 1993). This eliminated the undesired effects, and allowed plants to be viable and fertile, though mild effects of increased CK were occasionally observed in the non-targeted organs as well. Plant overexpressing *CKX* have been shown to contain reduced levels of cytokinin many times, in various plant species. *CKX3* overexpression was specifically shown to cause reduction in CKs in several works (Nishiyama et al., 2011; Reid et al., 2016). Reducing cytokinin levels with *CKX* overexpression led to stunting in Arabidopsis when expressed from the strong 35S promoter (Vercruyssen et al., 2011), but overexpression of *CKX3* in tomato had relatively minimal phenotypes under optimal conditions (Farber et al., 2016). The lines we used, which overexpress Arabidopsis *AtIPT7* or *AtCKX3* from the leaf specific promoters *pFIL* and *pBLS*, in tomato cv M82, have normal early development and are viable (Shani et al., 2010). *pFIL>>IPT7* is mostly infertile, which is why we used *pBLS>>IPT7*, that has a milder phenotype, due to the expression being later in development, and normal fertility (Shani et al., 2010). *pFIL>>CKX3* and *pFIL>>GFP-PROΔ17* are both viable and fertile. We used lines with expression driven from the *FIL* promoter in both these cases because lines driven from the *BLS* promoter had mild to undetectable leaf phenotypes. Thus, *BLS* was used only in the case of IPT, to avoid pleiotropic effects. CK and GA content were previously analyzed in the tomato *clausa* mutant. *clausa* is highly CK sensitive and GA insensitive, and displays meristematic and leaf phenotypes similar to those of overexpression of *IPT*. In terms of hormonal content, *clausa* has a significant reduction in the content of many CK compounds, and a significant increase in several active GA precursors (Israeli et al., 2021).

### Torin2 and WYE132 treatments

It was previously demonstrated that Torin2 and WYE132 are effective and specific inhibitors of TOR (Li et al., 2017; Marash et al., 2022; Montané & Menand, 2013). Torin2 (SML1224 Sigma-Aldrich) or WYE132 (PZ0321 Sigma-Aldrich), were applied to detached tomato leaves through the petiole for 24 hours prior to pathogen inoculation (both inhibitors), defense response quantification (only Torin2), or RNA preparation (only Torin2). For both inhibitors, a 10 mM stock solution was prepared in concentrated DMSO (P0037 SIGMA-Aldrich) and diluted to 2 µM in water. Mock leaves were treated with water containing 1:5000 of DMSO. Torin2 was used this study in concentration of 2 µM based on previous studies (Marash et al., 2022; Ye et al., 2022).

### Hormone treatments

For CK treatment (Figure S4), plants were sprayed with 100 μM 6-benzyl purine (6-BAP, Sigma-Aldrich) or Mock solution 24h prior to analysis. The stock solution was prepared in NaOH and diluted with water. Similarly diluted NaOH in water served as Mock.

For GA treatment (Figure 4), GA_3_ (Sigma-Aldrich) in the indicated concentrations was dissolved in ethanol and applied by spraying three times a week for two weeks. The stock solution was prepared in ethanol and diluted with water.

For assessing TOR activity following hormonal treatment (Figure 6), 100 μM 6-BAP or GA_3_, or both, were sprayed on developing leaves or mature L5, 4h prior to protein extraction. Mock treatments were dilute NaOH for CK, or ethanol with Tween 20 (100 μl/L) for GA, as described above.

### Virus Induced Gene Silencing (VIGS)

VIGS was performed as previously described (Liu *et al*., 2002). The *SlTOR* silencing construct was generated as previously described in Marash *et al*., 2022. The TRV2:TOR construct, as well as an empty TRV RNA2 for control, and the pTRV1 vector were introduced into *A. tumefaciens* strain GV3101::pMP90. The cultures were adjusted to OD_600_= 0.2 and TRV RNA1 was mixed in at a ratio of 1:1 with RNA2 (either empty or *TRV2:TOR*) in infiltration buffer, and infiltrated into cotyledons of 10-day-old seedlings. Analyses were subsequently conducted when plants reached 6 weeks of age.

### Pathogenesis assays

*Botrytis cinerea* (*Bc*) pathogenicity assays were performed as previously described (Gupta et al., 2020b). Briefly, inoculum of *Bc* isolate Bcl16 was maintained on potato dextrose agar (PDA; Difco) plates in an incubator at 22°C. Agar discs with a diameter of 0.4 cm were then pierced from colony margins and used to inoculate detached leaves. Inoculated leaves were kept in a humid chamber at 22°C under long-day conditions. Necrotic lesion size was measured 2-3 days post inoculation using ImageJ. For *Bc* inoculation prior to RT-qPCR analysis, *Bc* spores were collected from 21-day-old culture in 5 ml of 1 mg/ml glucose and 1 mg/ml K_2_HPO_4_. The spore suspension was vortexed, filtered through cheesecloth to remove mycelial fragments and used to inoculate tomato leaves by spraying. Mock plants were sprayed with similar concentrations of glucose and K_2_HPO_4_. RNA was prepared 24 h after *Bc* inoculation.

*Xanthomonas campestris pv. vesicatoria (Xcv)* was grown at 28°C in Luria Bertani (LB) broth overnight supplemented with Rifampicin (10 μg/ml) (Sigma), diluted to concentration of OD_600_=0.0002 in 10mM MgCl_2_, and used to pressure-infiltrate L5 of silenced plants with a 1-ml needleless syringe. Disease was assessed by measuring the water-soaked lesion area as previously described (Teper et al., 2018), 10 days after inoculation.

*Oidium neolycopersici (On)* was continuously maintained by randomly placing healthy plants alongside pre-infected plants in a dedicated chamber, and allowing them to be inoculated through air circulation. New plants were introduced every two weeks, upon which wholly covered chlorotic plants were discarded. For inoculation, diseased leaves with 80% On coverage were shaken above L5 of each plant. Two infected leaves were used for each individual. Disease severity was calculated based on the percentage of the leaf surface covered with powdery mildew symptoms 10-14 dpi.

### ROS production measurement

ROS measurement was carried out as previously described by (Leibman-Markus et al., 2017). 0.5 cm diameter leaf discs were collected, and each disc was incubated in 250µL distilled water in a 96-well plate (SPL Life Science) at room temperature with gentle shaking. After 4 hours, the water was removed and 50µL of distilled water were added. Right before measurement, 100µL of distilled water with or without 1µM flg22 (PhytoTechLabs #P6622) were added. Light emission was measured using a luminometer (GloMax® Discover, Promega, United States).

### Ion leakage (conductivity) measurement

Conductivity was measured according to (Leibman-Markus et al., 2017). 0.9 cm diameter leaf discs were harvested and washed with distilled water for 3 hours in a 50 mL tube. For each sample, five discs were placed in a 10-flask with 1 ml of distilled water, with 2µM Torin2, 6-BAP, or DMSO, for 48 hours at room temperature with gentle shaking. After incubation, 1.5 mL of distilled water were added to each sample, and conductivity was measured using a conductivity meter (AZ^®^ Multiparameter pH/Mv/Cond./Temp Meter 86505, Taiwan).

### RNA extraction and RT-qPCR

for RNA extraction, either five 0.9 cm diameter leaf discs were harvested from L5 of 6-week-old plants (Figure 5B, D, F, H) or whole shoots of of developing leaves (m+6) of 3-week-old seedlings with four true leaves (Figure 5A, C, E, G) were used. Isolation of total RNA was performed according to the TRI reagent (Sigma-Aldrich) procedure, with application of DNAse (EN0521 ThermoFisher) to remove genomic DNA. 1 µg of RNA was used for cDNA synthesis using Maxima reverse transcriptase (ThermoFisher). RT-qPCR assays were conducted with Power SYBR Green Mix (Life Technologies), using specific primers (Supplemental Table 1) in a Rotor-Gene Q machine (Qiagen). Standard curves were achieved by dilutions of one cDNA sample. Relative expression was quantified by dividing the expression of the relevant gene by the geometric mean of the expression of the following normalizer genes: for developing leaves: *RPL8* (Solyc10g006580), *EXP* (Solyc07g025390), and *CYP* (Solyc01g111170), and for mature leaves, *RPL8*, *CYP*, and *Actin* (Solyc11g005330). All primer pairs had efficiencies in the range of 0.97-1.03. All the primers used for RT-qPCR are listed in Table S1.

### Protein purification and Western Blotting

For protein purification from mature leaves, whole tomato leaves (Figures 6C-D and 7) were harvested from 6-week-old plants with nine true leaves, and ground in liquid nitrogen. 150 mg ground tissue was used. For developing leaves (Figure 6A,B), 25mg of tissue was collected from shoots of developing leaves (m+6) of 3-week-old seedling with four true leaves. Each sample of developing leaves was composed from a total of 10-12 different plants. The tissues were then ground in liquid nitrogen with 3 volumes of extraction buffer (100mM MOPS pH 7.6, 100mM NaCl, 40mM ß-MeOH, 5% SDS, 10% Glycerol, 4mM EDTA, 2mM PMSF and Pphosphatase Inhibitor (Sigma)), boiled for 5 minutes at 95°C and centrifuged at 10,000 rpm for 10 minutes, to remove cell debris. Samples of equal volume were separated on 15% SDS acrylamide gels, transferred to nitrocellulose membranes (Protran, #10401380), stained with Ponceau red as loading and transfer control, and blocked with 3% skimmed milk in Tris buffered saline (TBS) with 1% Tween20 for one hour at room temperature with gentle shaking. Membranes were probed with Anti-S6K1 p-Thr449 polyclonal antibody (AB-ab207399, Abcam,1:750), Anti-S6K1/2 (GRS-AS121855, Agrisera, 1:500), or Anti-ACTIN (GRS-AS132640, Agrisera, 1:1000) overnight at 4°C. IgG HRP-conjugated goat-anti-rabbit (AB-ab205718, Abcam, 1:10,000) was used as secondary antibody. Chemiluminescence was observed using Elistar Supernova as substrate (Cyanagen, #XLSE2) and images of protein bands were acquired and quantified using the Alliance UVITEC software. Phosphorylation status of S6K1 was tested 4 h after GA and CK treatment based on a previous study quantifying the decrease in TOR activity upon inhibition (Upadhyaya et al., 2020).

## Statistical analysis

All data are presented as average ±SEM, or as boxplots showing minimum to maximum values, with the box representing inner quartile ranges and the whiskers representing outer quartile ranges. Data sets were analyzed for normality using the Shapiro-Wilk test. For non-Gausian distributed samples, differences between two groups were analyzed for statistical significance using a Mann-Whitney U test, and differences between three groups or more were analyzed using Kruskal-Wallis ANOVA with Dunn’s post hoc test. For normally distributed samples, differences between two groups were analyzed for statistical significance using a two tailed t-test, with Welch’s correction for samples with unequal variances, where appropriate. Differences among three groups or more were analyzed for statistical significance using one-way ANOVA. Regular ANOVA was used for groups with equal variances, and Welch’s ANOVA for groups with unequal variances. When a significant result for a group in an ANOVA was returned, significance in differences between the means of different samples in the group was assessed using a post-hoc test. Tukey’s or Bonferroni’s tests were employed for samples with equal variances, and Dunnett’s test was employed for samples with unequal variances. All statistical analyses were conducted using Prism9^TM^.

## Data availability Statement

The authors declare that the data supporting the findings of this study are available within the paper and its Supplementary information files. Raw data is available from the corresponding author upon reasonable request.

## Supplementary data

**Figure S1:** TOR inhibition mediated disease resistance depends on the CK/GA balance-additional results.

**Figure S2:** *TOR* silencing mediated disease resistance depends on the CK/GA balance-VIGS results.

**Figure S3:** TOR inhibition mediated disease resistance depends on the CK/GA balance-ROS kinetics.

**Figure S4:** TOR inhibition mediated disease resistance to *X. campestris p. vesicatoria* and *O. neolycopersici* depends on the CK/GA balance.

**Figure S5:** TOR is required for the execution of developmental cues in the leaf.

**Figure S6:** TOR inhibition does not augment CK-mediated disease resistance.

**Figure S7:** *TOR* silencing by VIGS reduces CK response in the meristem and developing leaf primordia.

**Figure S8:** TOR inhibition and *Bc* infection have differential effects on CK pathway genes.

**Table S1:** Primer pairs used in this work.

## Notes

### Competing Interest Statement

The authors have declared no competing interest.

### Summary of Updates

Data from experiments concerning the relationship between Gibberellin, Cytokinin, and TOR was added to the revised manuscript.

## Bibliography

Achard, P., Cheng, H., De Grauwe, L., Decat, J., Schoutteten, H., Moritz, T., Van Der Straeten, D., Peng, J., & Harberd, N. P. (2006). Integration of plant responses to environmentally activated phytohormonal signals. Science, 311(5757), 91–94. https://doi.org/10.1126/science.1118642

Achard, P., Renou, J. P., Berthomé, R., Harberd, N. P., & Genschik, P. (2008). Plant DELLAs Restrain Growth and Promote Survival of Adversity by Reducing the Levels of Reactive Oxygen Species. Current Biology, 18(9), 656–660. https://doi.org/10.1016/j.cub.2008.04.034

Anderson, G. H., Veit, B., & Hanson, M. R. (2005). The Arabidopsis AtRaptor genes are essential for post-embryonic plant growth. BMC Biology, 3. https://doi.org/10.1186/1741-7007-3-12

Argueso, C. T., Ferreira, F. J., Epple, P., To, J. P. C., Hutchison, C. E., Schaller, G. E., Dangl, J. L., & Kieber, J. J. (2012). Two-component elements mediate interactions between cytokinin and salicylic acid in plant immunity. PLoS Genetics, 8(1). https://doi.org/10.1371/journal.pgen.1002448

Argyros, R. D., Mathews, D. E., Chiang, Y. H., Palmer, C. M., Thibault, D. M., Etheridge, N., Argyros, D. A., Mason, M. G., Kieber, J. J., & Schallera, G. E. (2008). Type B response regulators of Arabidopsis play key roles in cytokinin signaling and plant development. Plant Cell, 20(8), 2102–2116. https://doi.org/10.1105/tpc.108.059584

Aznar, N. R., Consolo, V. F., Salerno, G. L., & Martínez-Noël, G. M. A. (2018). TOR signaling downregulation increases resistance to the cereal killer Fusarium graminearum. Plant Signaling and Behavior, 13(2), e1414120. https://doi.org/10.1080/15592324.2017.1414120

Bar, M., Israeli, A., Levy, M., Gera, H. Ben, Jiménez-Gómez, J. M., Kouril, S., Tarkowski, P., & Ori, N. (2016). CLAUSA is a MYB transcription factor that promotes leaf differentiation by attenuating cytokinin signaling. Plant Cell, 28(7), 1602–1615. https://doi.org/10.1105/tpc.16.00211

Bari, R., & Jones, J. D. G. (2009). Role of plant hormones in plant defence responses. Plant Molecular Biology, 69(4), 473–488. https://doi.org/10.1007/s11103-008-9435-0

Bartrina, I., Otto, E., Strnad, M., Werner, T., & Schmülling, T. (2011). Cytokinin regulates the activity of reproductive meristems, flower organ size, ovule formation, and thus seed yield in Arabidopsis thaliana. Plant Cell, 23(1), 69–80. https://doi.org/10.1105/tpc.110.079079

Berens, M. L., Wolinska, K. W., Spaepen, S., Ziegler, J., Nobori, T., Nair, A., Krüler, V., Winkelmüller, T. M., Wang, Y., Mine, A., Becker, D., Garrido-Oter, R., Schulze-Lefert, P., & Tsuda, K. (2019). Balancing trade-offs between biotic and abiotic stress responses through leaf age-dependent variation in stress hormone cross-talk. Proceedings of the National Academy of Sciences of the United States of America, 116(6), 2364–2373. https://doi.org/10.1073/pnas.1817233116

Berry, H. M., & Argueso, C. T. (2022). More than growth: Phytohormone-regulated transcription factors controlling plant immunity, plant development and plant architecture. Current Opinion in Plant Biology, 70, 102309. https://doi.org/10.1016/j.pbi.2022.102309

Bolduc, N., & Hake, S. (2009). The maize transcription factor KNOTTED1 directly regulates the gibberellin catabolism gene ga2ox1. Plant Cell, 21(6), 1647–1658. https://doi.org/10.1105/tpc.109.068221

Brenner, W. G., Romanov, G. A., Köllmer, I., Bürkle, L., & Schmülling, T. (2005). Immediate-early and delayed cytokinin response genes of Arabidopsis thaliana identified by genome-wide expression profiling reveal novel cytokinin-sensitive processes and suggest cytokinin action through transcriptional cascades. Plant Journal, 44(2), 314–333. https://doi.org/10.1111/j.1365-313X.2005.02530.x

Brenner, W. G., & Schmülling, T. (2012). Transcript profiling of cytokinin action in Arabidopsis roots and shoots discovers largely similar but also organ-specific responses. BMC Plant Biology, 12. https://doi.org/10.1186/1471-2229-12-112

Brunkard, J. O., Xu, M., Regina Scarpin, M., Chatterjee, S., Shemyakina, E. A., Goodman, H. M., & Zambryski, P. (2020). TOR dynamically regulates plant cell-cell transport. Proceedings of the National Academy of Sciences of the United States of America, 117(9), 5049–5058. https://doi.org/10.1073/pnas.1919196117

Busch, B. L., Schmitz, G., Rossmann, S., Piron, F., Ding, J., Bendahmane, A., & Theres, K. (2011). Shoot branching and leaf dissection in tomato are regulated by homologous gene modules. Plant Cell, 23(10), 3595–3609. https://doi.org/10.1105/tpc.111.087981

Caldana, C., Li, Y., Leisse, A., Zhang, Y., Bartholomaeus, L., Fernie, A. R., Willmitzer, L., & Giavalisco, P. (2013). Systemic analysis of inducible target of rapamycin mutants reveal a general metabolic switch controlling growth in Arabidopsis thaliana. Plant Journal, 73(6), 897–909. https://doi.org/10.1111/tpj.12080

Caldana, C., Martins, M. C. M., Mubeen, U., & Castellanos, R. U. (2019). The magic ‘hammer’ of TOR: The multiple faces of a single pathway in the metabolic regulation of plant growth and development. Journal of Experimental Botany, 70(8), 2217–2225. https://doi.org/10.1093/jxb/ery459

Campos, M. L., Yoshida, Y., Major, I. T., De Oliveira Ferreira, D., Weraduwage, S. M., Froehlich, J. E., Johnson, B. F., Kramer, D. M., Jander, G., Sharkey, T. D., & Howe, G. A. (2016). Rewiring of jasmonate and phytochrome B signalling uncouples plant growth-defense tradeoffs. Nature Communications, 7. https://doi.org/10.1038/ncomms12570

Cao, P., Kim, S. J., Xing, A., Schenck, C. A., Liu, L., Jiang, N., Wang, J., Last, R. L., & Brandizzi, F. (2019). Homeostasis of branched-chain amino acids is critical for the activity of TOR signaling in Arabidopsis. ELife, 8. https://doi.org/10.7554/eLife.50747

Černý, M., Kuklová, A., Hoehenwarter, W., Fragner, L., Novák, O., Rotková, G., Jedelský, P. L., Žáková, K., Šmehilová, M., Strnad, M., Weckwerth, W., & Brzobohatý, B. (2013). Proteome and metabolome profiling of cytokinin action in Arabidopsis identifying both distinct and similar responses to cytokinin down- and up-regulation. Journal of Experimental Botany, 64(14), 4193–4206. https://doi.org/10.1093/jxb/ert227

Choi, J., Huh, S. U., Kojima, M., Sakakibara, H., Paek, K. H., & Hwang, I. (2010). The cytokinin-activated transcription factor ARR2 promotes plant immunity via TGA3/NPR1-dependent salicylic acid signaling in arabidopsis. Developmental Cell, 19(2), 284–295. https://doi.org/10.1016/j.devcel.2010.07.011

Colebrook, E. H., Thomas, S. G., Phillips, A. L., & Hedden, P. (2014). The role of gibberellin signalling in plant responses to abiotic stress. Journal of Experimental Biology, 217(1), 67–75. https://doi.org/10.1242/jeb.089938

Davière, J. M., & Achard, P. (2013). Gibberellin signaling in plants. Development (Cambridge), 140(6), 1147–1151. https://doi.org/10.1242/dev.087650

De Bruyne, L., Höfte, M., & De Vleesschauwer, D. (2014). Connecting growth and defense: The emerging roles of brassinosteroids and gibberellins in plant innate immunity. Molecular Plant, 7(6), 943–959. https://doi.org/10.1093/mp/ssu050

De Vleesschauwer, D., Filipe, O., Hoffman, G., Seifi, H. S., Haeck, A., Canlas, P., Van Bockhaven, J., De Waele, E., Demeestere, K., Ronald, P., & Hofte, M. (2018). Target of rapamycin signaling orchestrates growth-defense trade-offs in plants. New Phytologist, 217(1), 305–319. https://doi.org/10.1111/nph.14785

de Vleesschauwer, D., Seifi, H. S., Filipe, O., Haeck, A., Huu, S. N., Demeestere, K., & Höfte, M. (2016). The DELLA protein SLR1 integrates and amplifies salicylic acid- and jasmonic acid-dependent innate immunity in rice. Plant Physiology, 170(3), 1831–1847. https://doi.org/10.1104/pp.15.01515

de Vleesschauwer, D., van Buyten, E., Satoh, K., Balidion, J., Mauleon, R., Choi, I. R., Vera-Cruz, C., Kikuchi, S., & Höfte, M. (2012). Brassinosteroids antagonize gibberellin- and salicylate-mediated root immunity in rice. Plant Physiology, 158(4), 1833–1846. https://doi.org/10.1104/pp.112.193672

Deprost, D., Yao, L., Sormani, R., Moreau, M., Leterreux, G., Bedu, M., Robaglia, C., & Meyer, C. (2007). The Arabidopsis TOR kinase links plant growth, yield, stress resistance and mRNA translation. EMBO Reports, 8(9), 864–870. https://doi.org/10.1038/sj.embor.7401043

Develey-Rivière, M. P., & Galiana, E. (2007). Resistance to pathogens and host developmental stage: A multifaceted relationship within the plant kingdom. New Phytologist, 175(3), 405–416. https://doi.org/10.1111/j.1469-8137.2007.02130.x

Dobrenel, T., Caldana, C., Hanson, J., Robaglia, C., Vincentz, M., Veit, B., & Meyer, C. (2016). TOR Signaling and Nutrient Sensing. Annual Review of Plant Biology, 67(1), 261–285. https://doi.org/10.1146/annurev-arplant-043014-114648

Dong, P., Xiong, F., Que, Y., Wang, K., Yu, L., Li, Z., & Ren, M. (2015). Expression profiling and functional analysis reveals that TOR is a key player in regulating photosynthesis and phytohormone signaling pathways in Arabidopsis. Frontiers in Plant Science, 6(september), 1–15. https://doi.org/10.3389/fpls.2015.00677

Du, F., Guan, C., & Jiao, Y. (2018). Molecular Mechanisms of Leaf Morphogenesis. In Molecular Plant (Vol. 11, Issue 9, pp. 1117–1134). Cell Press. https://doi.org/10.1016/j.molp.2018.06.006

Efroni, I., Han, S. K., Kim, H. J., Wu, M. F., Steiner, E., Birnbaum, K. D., Hong, J. C., Eshed, Y., & Wagner, D. (2013). Regulation of leaf maturation by chromatin-mediated modulation of cytokinin responses. Developmental Cell, 24(4), 438–445. https://doi.org/10.1016/j.devcel.2013.01.019

Eichmann, R., & Schäfer, P. (2015). Growth versus immunity-a redirection of the cell cycle? Current Opinion in Plant Biology, 26, 106–112. https://doi.org/10.1016/j.pbi.2015.06.006

Ezura, H., & Harberd, N. P. (1995). Endogenous gibberellin levels influence in-vitro shoot regeneration in Arabidopsis thaliana (L.) Heynh. Planta, 197(2), 301–305. https://doi.org/10.1007/BF00202651

Farber, M., Attia, Z., & Weiss, D. (2016). Cytokinin activity increases stomatal density and transpiration rate in tomato. Journal of Experimental Botany, 67(22), 6351–6362. https://doi.org/10.1093/jxb/erw398

Fleishon, S., Shani, E., Ori, N., & Weiss, D. (2011). Negative reciprocal interactions between gibberellin and cytokinin in tomato. New Phytologist, 190(3), 609–617. https://doi.org/10.1111/j.1469-8137.2010.03616.x

Gan, S., & Amasino, R. M. (1995). Inhibition of leaf senescence by autoregulated production of cytokinin. Science, 270(5244), 1986–1988. https://doi.org/10.1126/science.270.5244.1986

Goss, E. M., & Bergelson, J. (2006). Variation in Resistance and Virulence in the Interaction Between Arabidopsis Thaliana and a Bacterial Pathogen. Evolution, 60(8), 1562. https://doi.org/10.1554/06-200.1

Greenboim-Wainberg, Y., Maymon, I., Borochov, R., Alvarez, J., Olszewski, N., Ori, N., Eshed, Y., & Weiss, D. (2005). Cross talk between gibberellin and cytokinin: The Arabidopsis GA response inhibitor SPINDLY plays a positive role in cytokinin signaling. Plant Cell, 17(1), 92–102. https://doi.org/10.1105/tpc.104.028472

Großkinsky, D. K., Naseem, M., Abdelmohsen, U. R., Plickert, N., Engelke, T., Griebel, T., Zeier, J., Novák, O., Strnad, M., Pfeifhofer, H., van der Graaff, E., Simon, U., & Roitsch, T. (2011). Cytokinins mediate resistance against Pseudomonas syringae in tobacco through increased antimicrobial phytoalexin synthesis independent of salicylic acid signaling. Plant Physiology, 157(2), 815–830. https://doi.org/10.1104/pp.111.182931

Gupta, R., Anand, G., Pizarro, L., Laor, D., Kovetz, N., Sela, N., Yehuda, T., Gazit, E., & Bar, M. (2021). Cytokinin Inhibits Fungal Development and Virulence by Targeting the Cytoskeleton and Cellular Trafficking. MBio, 12(5), 1–22. https://doi.org/10.1128/mBio.03068-20

Gupta, R., Elkabetz, D., Leibman-Markus, M., Jami, E., & Bar, M. (2022). Cytokinin-microbiome interactions regulate developmental functions. Environmental Microbiomes, 17(1), 1–16. https://doi.org/10.1186/s40793-022-00397-2

Gupta, R., Leibman-Markus, M., Pizarro, L., & Bar, M. (2021). Cytokinin induces bacterial pathogen resistance in tomato. Plant Pathology, 70(2), 318–325. https://doi.org/10.1111/ppa.13279

Gupta, R., Pizarro, L., Leibman-Markus, M., Marash, I., & Bar, M. (2020). Cytokinin response induces immunity and fungal pathogen resistance, and modulates trafficking of the PRR LeEIX2 in tomato. Molecular Plant Pathology, 10.1111/mpp.12978.

Harberd, N. P., Belfield, E., & Yasumura, Y. (2009). The angiosperm gibberellin-GID1-DELLA growth regulatory mechanism: How an “inhibitor of an inhibitor” enables flexible response to fluctuating environments. Plant Cell, 21(5), 1328–1339. https://doi.org/10.1105/tpc.109.066969

Hauvermale, A. L., Ariizumi, T., & Steber, C. M. (2012). Gibberellin signaling: A theme and variations on DELLA repression. Plant Physiology, 160(1), 83–92. https://doi.org/10.1104/pp.112.200956

Hay, A., Craft, J., & Tsiantis, M. (2004). Plant hormones and homeoboxes: Bridging the gap? BioEssays, 26(4), 395–404. https://doi.org/10.1002/bies.20016

Hay, A., Kaur, H., Phillips, A., Hedden, P., Hake, S., Tsiantis, M., Yanai, O., Shani, E., Dolezal, K., Tarkowski, P., Sablowski, R., Sandberg, G., Samach, A., & Ori, N. (2005). The gibberellin pathway mediates KNOTTED1-type homeobox function in plants with different body plans. Current Biology, 12(18), 1566–1571. https://doi.org/10.1016/j.cub.2005.07.060

Hay, A., & Tsiantis, M. (2010). KNOX genes: Versatile regulators of plant development and diversity. Development, 137(19), 3153–3165. https://doi.org/10.1242/dev.030049

Hedden, P. (2020). The current status of research on gibberellin biosynthesis. Plant and Cell Physiology, 61(11), 1832–1849. https://doi.org/10.1093/pcp/pcaa092

Horiguchi, G., Mollá-Morales, A., Pérez-Pérez, J. M., Kojima, K., Robles, P., Ponce, M. R., Micol, J. L., & Tsukaya, H. (2011). Differential contributions of ribosomal protein genes to arabidopsis thaliana leaf development. Plant Journal, 65(5), 724–736. https://doi.org/10.1111/j.1365-313X.2010.04457.x

Hou, X., Lee, L. Y. C., Xia, K., Yan, Y., & Yu, H. (2010). DELLAs Modulate Jasmonate Signaling via Competitive Binding to JAZs. Developmental Cell, 19(6), 884–894. https://doi.org/10.1016/j.devcel.2010.10.024

Hwang, I., Sheen, J., & Müller, B. (2012). Cytokinin signaling networks. In Annual Review of Plant Biology (Vol. 63, pp. 353–380). https://doi.org/10.1146/annurev-arplant-042811-105503

Hwang, I., Sheen, J., Müller, B., Riefler, M., Novak, O., Strnad, M., Schmülling, T., Kim, H. J., Ryu, H., Hong, S. H., Woo, H. R., Lim, P. O., Lee, I. C., Sheen, J., Nam, H. G., Hwang, I., Gan, S., & Amasino, R. M. (2006). Cytokinin-mediated control of leaf longevity by AHK3 through phosphorylation of ARR2 in Arabidopsis. Plant Cell, 18(1), 1986–1988. https://doi.org/10.1126/science.270.5244.1986

Ishida, K., Yamashino, T., Yokoyama, A., & Mizuno, T. (2008). Three type-B response regulators, ARR1, ARR10 and ARR12, play essential but redundant roles in cytokinin signal transduction throughout the life cycle of Arabidopsis thaliana. Plant and Cell Physiology, 49(1), 47–57. https://doi.org/10.1093/pcp/pcm165

Israeli, A., Burko, Y., Shleizer-Burko, S., Zelnik, I. D., Sela, N., Hajirezaei, M. R., Fernie, A. R., Tohge, T., Ori, N., & Bar, M. (2021). Coordinating the morphogenesis-differentiation balance by tweaking the cytokinin-gibberellin equilibrium. PLoS Genetics, 17(4), 1–25. https://doi.org/10.1371/journal.pgen.1009537

Israeli, A., Capua, Y., Shwartz, I., Tal, L., Meir, Z., Levy, M., Bar, M., Efroni, I., & Ori, N. (2019). Multiple Auxin-Response Regulators Enable Stability and Variability in Leaf Development. Current Biology, 29(11), 1746–1759.e5. https://doi.org/10.1016/j.cub.2019.04.047

Jacob, D., Rav David, D., Sztjenberg, A., & Elad, Y. (2008). Conditions for development of powdery mildew of tomato caused by Oidium neolycopersici. Phytopathology, 98(3), 270–281. https://doi.org/10.1094/PHYTO-98-3-0270

Jasinski, S., Piazza, P., Craft, J., Hay, A., Woolley, L., Rieu, I., Phillips, A., Hedden, P., & Tsiantis, M. (2005). KNOX action in Arabidopsis is mediated by coordinate regulation of cytokinin and gibberellin activities. Current Biology, 15(17), 1560–1565. https://doi.org/10.1016/j.cub.2005.07.023

Jasinski, S., Tattersall, A., Piazza, P., Hay, A., Martinez-garcia, J. F., Schmitz, G., Theres, K., Mccormick, S., & Tsiantis, M. (2008). PROCERA encodes a DELLA protein that mediates control of dissected leaf form in tomato. 603–612. https://doi.org/10.1111/j.1365-313X.2008.03628.x

Jiang, C., Fan, Z., Li, Z., Niu, D., Li, Y., Zheng, M., Wang, Q., Jin, H., & Guo, J. (2020). Bacillus cereus AR156 triggers induced systemic resistance against Pseudomonas syringae pv. tomato DC3000 by suppressing miR472 and activating CNLs-mediated basal immunity in Arabidopsis. Molecular Plant Pathology, 21(6), 854–870. https://doi.org/10.1111/mpp.12935

Jiang, C. J., Shimono, M., Sugano, S., Kojima, M., Liu, X., Inoue, H., Sakakibara, H., & Takatsuji, H. (2013). Cytokinins act synergistically with salicylic acid to activate defense gene expression in rice. Molecular Plant-Microbe Interactions, 26(3), 287–296. https://doi.org/10.1094/MPMI-06-12-0152-R

Kakimoto, T. (2001). Identification of plant cytokinin biosynthetic enzymes as dimethylallyl diphosphate:ATP/ADP isopentenyltransferases. Plant and Cell Physiology, 42(7), 677–685. https://doi.org/10.1093/pcp/pce112

Karasov, T. L., Chae, E., Herman, J. J., & Bergelson, J. (2017). Mechanisms to mitigate the trade-off between growth and defense. Plant Cell, 29(4), 666–680. https://doi.org/10.1105/tpc.16.00931

Kazan, K., & Manners, J. M. (2013). MYC2: The master in action. Molecular Plant, 6(3), 686–703. https://doi.org/10.1093/mp/sss128

Kimura, S., Koenig, D., Kang, J., Yoong, F. Y., & Sinha, N. (2008). Natural Variation in Leaf Morphology Results from Mutation of a Novel KNOX Gene. Current Biology, 18(9), 672–677. https://doi.org/10.1016/j.cub.2008.04.008

Kliebenstein, D. J. (2016). False idolatry of the mythical growth versus immunity tradeoff in molecular systems plant pathology. Physiological and Molecular Plant Pathology, 95, 55–59. https://doi.org/10.1016/j.pmpp.2016.02.004

Kurakawa, T., Ueda, N., Maekawa, M., Kobayashi, K., Kojima, M., Nagato, Y., Sakakibara, H., & Kyozuka, J. (2007). Direct control of shoot meristem activity by a cytokinin-activating enzyme. Nature, 445(7128), 652–655. https://doi.org/10.1038/nature05504

Kuroha, T., Tokunaga, H., Kojima, M., Ueda, N., Ishida, T., Nagawa, S., Fukuda, H., Sugimoto, K., & Sakakibara, H. (2009). Functional analyses of LONELY GUY cytokinin-activating enzymes reveal the importance of the direct activation pathway in Arabidopsis. Plant Cell, 21(10), 3152–3169. https://doi.org/10.1105/tpc.109.068676

Li, X., Cai, W., Liu, Y., Li, H., Fu, L., Liu, Z., Xu, L., Liu, H., Xu, T., & Xiong, Y. (2017). Differential TOR activation and cell proliferation in Arabidopsis root and shoot apexes. Proceedings of the National Academy of Sciences of the United States of America, 114(10), 2765–2770. https://doi.org/10.1073/pnas.1618782114

Lifschitz, E., Eviatar, T., Rozman, A., Shalit, A., Goldshmidt, A., Amsellem, Z., Alvarez, J. P., & Eshed, Y. (2006). The tomato FT ortholog triggers systemic signals that regulate growth and flowering and substitute for diverse environmental stimuli. Proceedings of the National Academy of Sciences of the United States of America, 103(16), 6398–6403. https://doi.org/10.1073/pnas.0601620103

Liu, Y., & Bassham, D. C. (2010). TOR is a negative regulator of autophagy in Arabidopsis thaliana. PLoS ONE, 5(7). https://doi.org/10.1371/journal.pone.0011883

Liu, Y., Duan, X., Zhao, X., Ding, W., Wang, Y., & Xiong, Y. (2021). Diverse nitrogen signals activate convergent ROP2-TOR signaling in Arabidopsis. Developmental Cell, 56(9), 1283–1295.e5. https://doi.org/10.1016/j.devcel.2021.03.022

Livne, S., Lor, V. S., Nir, I., Eliaz, N., Aharoni, A., Olszewski, N. E., Eshed, Y., & Weiss, D. (2015). Uncovering DELLA-independent gibberellin responses by characterizing new tomato procera mutants. Plant Cell, 27(6), 1579–1594. https://doi.org/10.1105/tpc.114.132795

Locascio, A., Blázquez, M. A., & Alabadí, D. (2013). Genomic analysis of della protein activity. Plant and Cell Physiology, 54(8), 1229–1237. https://doi.org/10.1093/pcp/pct082

Major, I. T., Guo, Q., Zhai, J., Kapali, G., Kramer, D. M., & Howea, G. A. (2020). A phytochrome b-independent pathway restricts growth at high levels of jasmonate defense. Plant Physiology, 183(2), 733–749. https://doi.org/10.1104/pp.19.01335

Mao, Y. B., Liu, Y. Q., Chen, D. Y., Chen, F. Y., Fang, X., Hong, G. J., Wang, L. J., Wang, J. W., & Chen, X. Y. (2017). Jasmonate response decay and defense metabolite accumulation contributes to age-regulated dynamics of plant insect resistance. Nature Communications, 8, 1–13. https://doi.org/10.1038/ncomms13925

Marash, I., Leibman-Markus, M., Gupta, R., Avni, A., & Bar, M. (2022). TOR inhibition primes immunity and pathogen resistance in tomato in a salicylic acid-dependent manner. Molecular Plant Pathology, 23(7), 1035–1047. https://doi.org/10.1111/mpp.13207

Margalha, L., Confraria, A., & Baena-González, E. (2019). SnRK1 and TOR: Modulating growth–defense trade-offs in plant stress responses. In Journal of Experimental Botany (Vol. 70, Issue 8, pp. 2261–2274). Oxford University Press. https://doi.org/10.1093/jxb/erz066

Márquez-López, R. E., Quintana-Escobar, A. O., & Loyola-Vargas, V. M. (2019). Cytokinins, the Cinderella of plant growth regulators. Phytochemistry Reviews, 18(6), 1387–1408. https://doi.org/10.1007/s11101-019-09656-6

McCready, K., Spencer, V., & Kim, M. (2020). The Importance of TOR Kinase in Plant Development. Frontiers in Plant Science, 11(February), 1–7. https://doi.org/10.3389/fpls.2020.00016

Meldau, S., Erb, M., & Baldwin, I. T. (2012). Defence on demand: Mechanisms behind optimal defence patterns. Annals of Botany, 110(8), 1503–1514. https://doi.org/10.1093/aob/mcs212

Meteignier, L. V., El Oirdi, M., Cohen, M., Barff, T., Matteau, D., Lucier, J. F., Rodrigue, S., Jacques, P. E., Yoshioka, K., & Moffett, P. (2018). Corrigendum: Translatome analysis of an NB-LRR immune response identifies important contributors to plant immunity in Arabidopsis (Journal of Experimental Botany (2017) 68:9 (2333-2344) DOI: 10.1093/jxb/erx078). In Journal of Experimental Botany (Vol. 69, Issue 15, p. 3785). Oxford University Press. https://doi.org/10.1093/jxb/ery150

Mok, D. W. S., & Mok, M. C. (2001). Etabolism and. Perception, 52(39), 89–118. http://www.ncbi.nlm.nih.gov/pubmed/11337393

Monson, R. K., Trowbridge, A. M., Lindroth, R. L., & Lerdau, M. T. (2022). Coordinated resource allocation to plant growth–defense tradeoffs. New Phytologist, 233(3), 1051–1066. https://doi.org/10.1111/nph.17773

Montané, M. H., & Menand, B. (2013). ATP-competitive mTOR kinase inhibitors delay plant growth by triggering early differentiation of meristematic cells but no developmental patterning change. Journal of Experimental Botany, 64(14), 4361–4374. https://doi.org/10.1093/jxb/ert242

Moreau, M., Azzopardi, M., Clé ment, G., Dobrenel, T., Marchive, C., Renne, C., Martin-Magniette, M. L., Taconnat, L., Renou, J. P., Robaglia, C., & Meyer, C. (2012). Mutations in the Arabidopsis homolog of LST8/GβL, a partner of the target of Rapamycin kinase, impair plant growth, flowering, and metabolic adaptation to long days. Plant Cell, 24(2), 463–481. https://doi.org/10.1105/tpc.111.091306

Moss, W. P., Byrne, J. M., Campbell, H. L., Ji, P., Bonas, U., Jones, J. B., & Wilson, M. (2007). Biological control of bacterial spot of tomato using hrp mutants of Xanthomonas campestris pv. vesicatoria. Biological Control, 41(2), 199–206. https://doi.org/10.1016/j.biocontrol.2007.01.008

Murase, K., Hirano, Y., Sun, T. P., & Hakoshima, T. (2008). Gibberellin-induced DELLA recognition by the gibberellin receptor GID1. Nature, 456(7221), 459–463. https://doi.org/10.1038/nature07519

Nakayama, H., Rowland, S. D., Cheng, Z., Zumstein, K., Kang, J., Kondo, Y., & Sinha, N. R. (2021). Leaf form diversification in an ornamental heirloom tomato results from alterations in two different HOMEOBOX genes. Current Biology, 31(21), 4788–4799.e5. https://doi.org/10.1016/j.cub.2021.08.023

Naseem, M., Philippi, N., Hussain, A., Wangorsch, G., Ahmed, N., & Dandekara, T. (2012). Integrated systems view on Networking by hormones in Arabidopsis immunity reveals multiple crosstalk for cytokinin. Plant Cell, 24(5), 1793–1814. https://doi.org/10.1105/tpc.112.098335

Navarro, L., Bari, R., Achard, P., Lisón, P., Nemri, A., Harberd, N. P., & Jones, J. D. G. (2008). DELLAs Control Plant Immune Responses by Modulating the Balance of Jasmonic Acid and Salicylic Acid Signaling. Current Biology, 18(9), 650–655. https://doi.org/10.1016/j.cub.2008.03.060

Nelissen, H., Rymen, B., Jikumaru, Y., Demuynck, K., Van Lijsebettens, M., Kamiya, Y., Inzé, D., & Beemster, G. T. S. (2012). A local maximum in gibberellin levels regulates maize leaf growth by spatial control of cell division. Current Biology, 22(13), 1183–1187. https://doi.org/10.1016/j.cub.2012.04.065

Nir, I., Shohat, H., Panizel, I., Olszewski, N., Aharoni, A., & Weiss, D. (2017). The tomato DELLA protein PROCERA acts in guard cells to promote stomatal closure. Plant Cell, 29(12), 3186–3197. https://doi.org/10.1105/tpc.17.00542

Nishiyama, R., Watanabe, Y., Fujita, Y., Le, D. T., Kojima, M., Werner, T., Vankova, R., Yamaguchi-Shinozaki, K., Shinozaki, K., Kakimoto, T., Sakakibara, H., Schmülling, T., & Tran, L. S. P. (2011). Analysis of cytokinin mutants and regulation of cytokinin metabolic genes reveals important regulatory roles of cytokinins in drought, salt and abscisic acid responses, and abscisic acid biosynthesis. Plant Cell, 23(6), 2169–2183. https://doi.org/10.1105/tpc.111.087395

Ori, N., Cohen, A. R., Etzioni, A., Brand, A., Yanai, O., Shleizer, S., Menda, N., Amsellem, Z., Efroni, I., Pekker, I., Alvarez, J. P., Blum, E., Zamir, D., & Eshed, Y. (2007). Regulation of LANCEOLATE by miR319 is required for compound-leaf development in tomato. Nature Genetics, 39(6), 787–791. https://doi.org/10.1038/ng2036

Pereyra, C. M., Aznar, N. R., Rodriguez, M. S., Salerno, G. L., & Martínez-Noël, G. M. A. (2020). Target of rapamycin signaling is tightly and differently regulated in the plant response under distinct abiotic stresses. Planta, 251(1), 1–14. https://doi.org/10.1007/s00425-019-03305-0

Porri, A., Torti, S., Mateos, J., Romera-Branchat, M., García-Martínez, J. L., Fornara, F., Gregis, V., Kater, M. M., & Coupland, G. (2014). SHORT VEGETATIVE PHASE reduces gibberellin biosynthesis at the Arabidopsis shoot apex to regulate the floral transition Fernando Andrés1. Proceedings of the National Academy of Sciences of the United States of America, 111(26). https://doi.org/10.1073/pnas.1409567111

Qin, X., Liu, J. H., Zhao, W. S., Chen, X. J., Guo, Z. J., & Peng, Y. L. (2013). Gibberellin 20-Oxidase Gene OsGA20ox3 regulates plant stature and disease development in rice. Molecular Plant-Microbe Interactions, 26(2), 227–239. https://doi.org/10.1094/MPMI-05-12-0138-R

Redig, P., Schmülling, T., & Van Onckelen, H. (1996). Analysis of cytokinin metabolism in ipt transgenic tobacco by liquid chromatography-tandem mass spectrometry. Plant Physiology, 112(1), 141–148. https://doi.org/10.1104/pp.112.1.141

Reid, D. E., Heckmann, A. B., Novák, O., Kelly, S., & Stougaard, J. (2016). CYTOKININ OXIDASE/DEHYDROGENASE3 maintains cytokinin homeostasis during root and nodule development in Lotus Japonicus. Plant Physiology, 170(2), 1060–1074. https://doi.org/10.1104/pp.15.00650

Riefler, M., Novak, O., Strnad, M., & Schmülling, T. (2006). Arabidopsis cytokinin receptors mutants reveal functions in shoot growth, leaf senescence, seed size, germination, root development, and cytokinin metabolism. Plant Cell, 18(1), 40–54. https://doi.org/10.1105/tpc.105.037796

Ryabova, L. A., Robaglia, C., & Meyer, C. (2019). Target of Rapamycin kinase: Central regulatory hub for plant growth and metabolism. Journal of Experimental Botany, 70(8), 2211–2216. https://doi.org/10.1093/jxb/erz108

Sakamoto, T., Kamiya, N., Ueguehi-Tanaka, M., Iwahori, S., & Matsuoka, M. (2001). KNOX homeodomain protein directly suppresses the expression of a gibberellin biosynthetic gene in the tobacco shoot apical meristem. Genes and Development, 15(5), 581–590. https://doi.org/10.1101/gad.867901

Sakamoto, T., Sakakibara, H., Kojima, M., Yamamoto, Y., Nagasaki, H., Inukai, Y., Sato, Y., & Matsuoka, M. (2006). Ectopic expression of KNOTTED1-like homeobox protein induces expression of cytokinin biosynthesis genes in rice. Plant Physiology, 142(1), 54–62. https://doi.org/10.1104/pp.106.085811

Sambrook, J., & Russell, D. W. (2006). SDS-Polyacrylamide Gel Electrophoresis of Proteins. Cold Spring Harbor Protocols, 2006(4), pdb.prot4540. https://doi.org/10.1101/pdb.prot4540

Saxton, R. A., & Sabatini, D. M. (2017). mTOR Signaling in Growth, Metabolism, and Disease. In Cell (Vol. 168, Issue 6, pp. 960–976). Cell Press. https://doi.org/10.1016/j.cell.2017.02.004

Shani, E., Ben-Gera, H., Shleizer-Burko, S., Burko, Y., Weiss, D., & Ori, N. (2010). Cytokinin regulates compound leaf development in tomato c w. Plant Cell, 22(10), 3206–3217. https://doi.org/10.1105/tpc.110.078253

Shani, E., Yanai, O., & Ori, N. (2006). The role of hormones in shoot apical meristem function. Current Opinion in Plant Biology, 9(5), 484–489. https://doi.org/10.1016/j.pbi.2006.07.008

Shi, L., & Olszewski, N. E. (1998). Gibberellin and abscisic acid regulate GAST1 expression at the level of transcription. Plant Molecular Biology, 38(6), 1053–1060. https://doi.org/10.1023/A:1006007315718

Shi, L., Wu, Y., & Sheen, J. (2018). TOR signaling in plants: conservation and innovation. In Development (Cambridge, England) (Vol. 145, Issue 13). https://doi.org/10.1242/dev.160887

Shwartz, I., Levy, M., Ori, N., & Bar, M. (2016). Hormones in tomato leaf development. Developmental Biology, 419(1), 132–142. https://doi.org/10.1016/j.ydbio.2016.06.023

Smailov, B., Alybayev, S., Smekenov, I., Mursalimov, A., Saparbaev, M., Sarbassov, D., & Bissenbaev, A. (2020). Wheat Germination Is Dependent on Plant Target of Rapamycin Signaling. Frontiers in Cell and Developmental Biology, 8(November). https://doi.org/10.3389/fcell.2020.606685

Smigocki, A. C., & Owens, L. D. (1988). Cytokinin gene fused with a strong promoter enhances shoot organogenesis and zeatin levels in transformed plant cells. Proceedings of the National Academy of Sciences, 85(14), 5131–5135. https://doi.org/10.1073/pnas.85.14.5131

Smigocki, A., Neal, J. W., McCanna, I., & Douglass, L. (1993). Cytokinin-mediated insect resistance in Nicotiana plants transformed with the ipt gene. Plant Molecular Biology, 23(2), 325–335. https://doi.org/10.1007/BF00029008

Song, L., Xu, G., Li, T., Zhou, H., Lin, Q., Chen, J., Wang, L., Wu, D., Li, X., Wang, L., Zhu, S., & Yu, F. (2022). The RALF1-FERONIA complex interacts with and activates TOR signaling in response to low nutrients. Molecular Plant, 15(7), 1120–1136. https://doi.org/10.1016/j.molp.2022.05.004

Song, Y., Zhao, G., Zhang, X., Li, L., Xiong, F., Zhuo, F., Zhang, C., Yang, Z., Datla, R., Ren, M., & Li, F. (2017). The crosstalk between Target of Rapamycin (TOR) and Jasmonic Acid (JA) signaling existing in Arabidopsis and cotton. Scientific Reports, 7(1), 1–15. https://doi.org/10.1038/srep45830

Soprano, A. S., Smetana, J. H. C., & Benedetti, C. E. (2018). Regulation of tRNA biogenesis in plants and its link to plant growth and response to pathogens. Biochimica et Biophysica Acta - Gene Regulatory Mechanisms, 1861(4), 344–353. https://doi.org/10.1016/j.bbagrm.2017.12.004

Steiner, E., Israeli, A., Gupta, R., Shwartz, I., Nir, I., Leibman-Markus, M., Tal, L., Farber, M., Amsalem, Z., Ori, N., Müller, B., & Bar, M. (2020). Characterization of the cytokinin sensor TCSv2 in arabidopsis and tomato. Plant Methods, 16(1), 1–12. https://doi.org/10.1186/s13007-020-00694-2

Steiner, E., Livne, S., Kobinson-Katz, T., Tal, L., Pri-Tal, O., Mosquna, A., Tarkowská, D., Mueller, B., Tarkowski, P., & Weiss, D. (2016). The putative O-linked N-acetylglucosamine transferase SPINDLY inhibits class I TCP proteolysis to promote sensitivity to cytokinin. Plant Physiology, 171(2), 1485–1494. https://doi.org/10.1104/pp.16.00343

Sun, T. ping. (2010). Gibberellin-GID1-DELLA: A pivotal regulatory module for plant growth and development. Plant Physiology, 154(2), 567–570. https://doi.org/10.1104/pp.110.161554

Sun, X., Jones, W. T., & Rikkerink, E. H. A. (2012). GRAS proteins: The versatile roles of intrinsically disordered proteins in plant signalling. Biochemical Journal, 442(1), 1–12. https://doi.org/10.1042/BJ20111766

Takei, K., Sakakibara, H., & Sugiyama, T. (2001). Identification of Genes Encoding Adenylate Isopentenyltransferase, a Cytokinin Biosynthesis Enzyme, in Arabidopsis thaliana. Journal of Biological Chemistry, 276(28), 26405–26410. https://doi.org/10.1074/jbc.M102130200

Teper, D., Girija, A. M., Bosis, E., Popov, G., Savidor, A., and Sessa, G. (2018). The Xanthomonas euvesicatoria type III effector XopAU is an active protein kinase that manipulates plant MAP kinase signaling. PLOS Pathogens 14, e1006880. doi: 10.1371/journal.ppat.1006880.

Turck, F., Zilbermann, F., Kozma, S. C., Thomas, G., & Nagy, F. (2004). Phytohormones participate in an S6 kinase signal transduction pathway in arabidopsis. Plant Physiology, 134(4), 1527–1535. https://doi.org/10.1104/pp.103.035873

Upadhyaya, S., Agrawal, S., Gorakshakar, A., & Rao, B. J. (2020). TOR kinase activity in Chlamydomonas reinhardtii is modulated by cellular metabolic states. FEBS Letters, 594(19), 3122–3141. https://doi.org/10.1002/1873-3468.13888

Vercruyssen, L., Gonzalez, N., Werner, T., Schmülling, T., & Inzé, D. (2011). Combining enhanced root and shoot growth reveals cross talk between pathways that control plant organ size in arabidopsis. Plant Physiology, 155(3), 1339–1352. https://doi.org/10.1104/pp.110.167049

Verma, R. K., & Teper, D. (2022). Immune recognition of the secreted serine protease ChpG restricts the host range of Clavibacter michiganensis from eggplant varieties. Molecular Plant Pathology, 23(7), 933–946. https://doi.org/10.1111/mpp.13215

Wang, P., Zhao, Y., Li, Z., Hsu, C. C., Liu, X., Fu, L., Hou, Y. J., Du, Y., Xie, S., Zhang, C., Gao, J., Cao, M., Huang, X., Zhu, Y., Tang, K., Wang, X., Tao, W. A., Xiong, Y., & Zhu, J. K. (2018). Reciprocal Regulation of the TOR Kinase and ABA Receptor Balances Plant Growth and Stress Response. Molecular Cell, 69(1), 100–112.e6. https://doi.org/10.1016/j.molcel.2017.12.002

Wang, X., Han, F., Yang, M., Yang, P., & Shen, S. (2013). Exploring the response of rice (Oryza sativa) leaf to gibberellins: A proteomic strategy. Rice, 6(1), 1–12. https://doi.org/10.1186/1939-8433-6-17

Weiss, D., & Ori, N. (2007). Mechanisms of cross talk between gibberellin and other hormones. Plant Physiology, 144(3), 1240–1246. https://doi.org/10.1104/pp.107.100370

Werner I; Bartrina, I; Holst, K; Schmülling, T, T. K. (2006). New Insights into the Biology of Cytokinin Degradation. Plant Biol (Stuttg), 8(03), 371–381. https://doi.org/10.1055/s-2006-923928

Werner, T., Motyka, V., Laucou, V., Smets, R., Van Onckelen, H., & Schmülling, T. (2003). Cytokinin-Deficient Transgenic Arabidopsis Plants Show Multiple Developmental Alterations Indicating Opposite Functions of Cytokinins in the Regulation of Shoot and Root Meristem Activity. Plant Cell, 15(11), 2532–2550. https://doi.org/10.1105/tpc.014928

Xiong, Y., McCormack, M., Li, L., Hall, Q., Xiang, C., & Sheen, J. (2013). Glucose-TOR signalling reprograms the transcriptome and activates meristems. Nature, 496(7444), 181–186. https://doi.org/10.1038/nature12030

Xiong, Y., & Sheen, J. (2012). Rapamycin and glucose-target of rapamycin (TOR) protein signaling in plants. Journal of Biological Chemistry, 287(4), 2836–2842. https://doi.org/10.1074/jbc.M111.300749

Xu, Y. P., Lv, L. H., Xu, Y. J., Yang, J., Cao, J. Y., & Cai, X. Z. (2018). Leaf stage-associated resistance is correlated with phytohormones in a pathosystem-dependent manner. Journal of Integrative Plant Biology, 60(8), 703–722. https://doi.org/10.1111/jipb.12661

Yan, J., Zhang, C., Gu, M., Bai, Z., Zhang, W., Qi, T., Cheng, Z., Peng, W., Luo, H., Nan, F., Wang, Z., & Xie, D. (2009). The arabidopsis CORONATINE INSENSITIVE1 protein is a jasmonate receptor. Plant Cell, 21(8), 2220–2236. https://doi.org/10.1105/tpc.109.065730

Yanai, O., Shani, E., Dolezal, K., Tarkowski, P., Sablowski, R., Sandberg, G., Samach, A., & Ori, N. (2005). Arabidopsis KNOXI proteins activate cytokinin biosynthesis. Current Biology, 15(17), 1566–1571. https://doi.org/10.1016/j.cub.2005.07.060

Yanai, O., Shani, E., Russ, D., & Ori, N. (2011). Gibberellin partly mediates LANCEOLATE activity in tomato. Plant Journal, 68(4), 571–582. https://doi.org/10.1111/j.1365-313X.2011.04716.x

Yang, D. L., Li, Q., Deng, Y. W., Lou, Y. G., Wang, M. Y., Zhou, G. X., Zhang, Y. Y., & He, Z. H. (2008). Altered disease development in the eui mutants and Eui overexpressors indicates that gibberellins negatively regulate rice basal disease resistance. Molecular Plant, 1(3), 528–537. https://doi.org/10.1093/mp/ssn021

Ye, R., Wang, M., Du, H., Chhajed, S., Koh, J., Liu, K. hsiang, Shin, J., Wu, Y., Shi, L., Xu, L., Chen, S., Zhang, Y., & Sheen, J. (2022). Glucose-driven TOR–FIE–PRC2 signalling controls plant development. Nature, 609(7929), 986–993. https://doi.org/10.1038/s41586-022-05171-5

Yuan, X., Xu, P., Yu, Y., & Xiong, Y. (2020). Glucose-TOR signaling regulates PIN2 stability to orchestrate auxin gradient and cell expansion in Arabidopsis root. Proceedings of the National Academy of Sciences of the United States of America, 117(51), 32223–32225. https://doi.org/10.1073/pnas.2015400117

Zeier, J. (2005). Age-dependent variations of local and systemic defence responses in Arabidopsis leaves towards an avirulent strain of Pseudomonas syringae. Physiological and Molecular Plant Pathology, 66(1–2), 30–39. https://doi.org/10.1016/j.pmpp.2005.03.007

Zhang, P., & Chen, K. (2009). Age-dependent variations of volatile emissions and inhibitory activity toward Botrytis cinerea and Fusarium oxysporum in tomato leaves treated with chitosan oligosaccharide. Journal of Plant Biology, 52(4), 332–339. https://doi.org/10.1007/s12374-009-9043-9

Zhang, Y., Zhang, Y., McFarlane, H. E., Obata, T., Richter, A. S., Lohse, M., Grimm, B., Persson, S., Fernie, A. R., & Giavalisco, P. (2018). Inhibition of TOR represses nutrient consumption, which improves greening after extended periods of etiolation. Plant Physiology, 178(1), 101–117. https://doi.org/10.1104/pp.18.00684

Zhang, Z., Zhu, J. Y., Roh, J., Marchive, C., Kim, S. K., Meyer, C., Sun, Y., Wang, W., & Wang, Z. Y. (2016). TOR Signaling Promotes Accumulation of BZR1 to Balance Growth with Carbon Availability in Arabidopsis. Current Biology, 26(14), 1854–1860. https://doi.org/10.1016/j.cub.2016.05.005

Zürcher, E., & Müller, B. (2016). Cytokinin Synthesis, Signaling, and Function-Advances and New Insights. In International Review of Cell and Molecular Biology (Vol. 324, pp. 1–38). Elsevier Inc. https://doi.org/10.1016/bs.ircmb.2016.01.001

Zürcher, E., Tavor-Deslex, D., Lituiev, D., Enkerli, K., Tarr, P. T., & Müller, B. (2013). A robust and sensitive synthetic sensor to monitor the transcriptional output of the cytokinin signaling network in planta. Plant Physiology, 161(3), 1066–1075. https://doi.org/10.1104/pp.112.211763

