## Supplementary material for "TOR coordinates Cytokinin and Gibberellin signals mediating development and defense": All Supplemental Figures and Table in one file

**Supplemental materials**

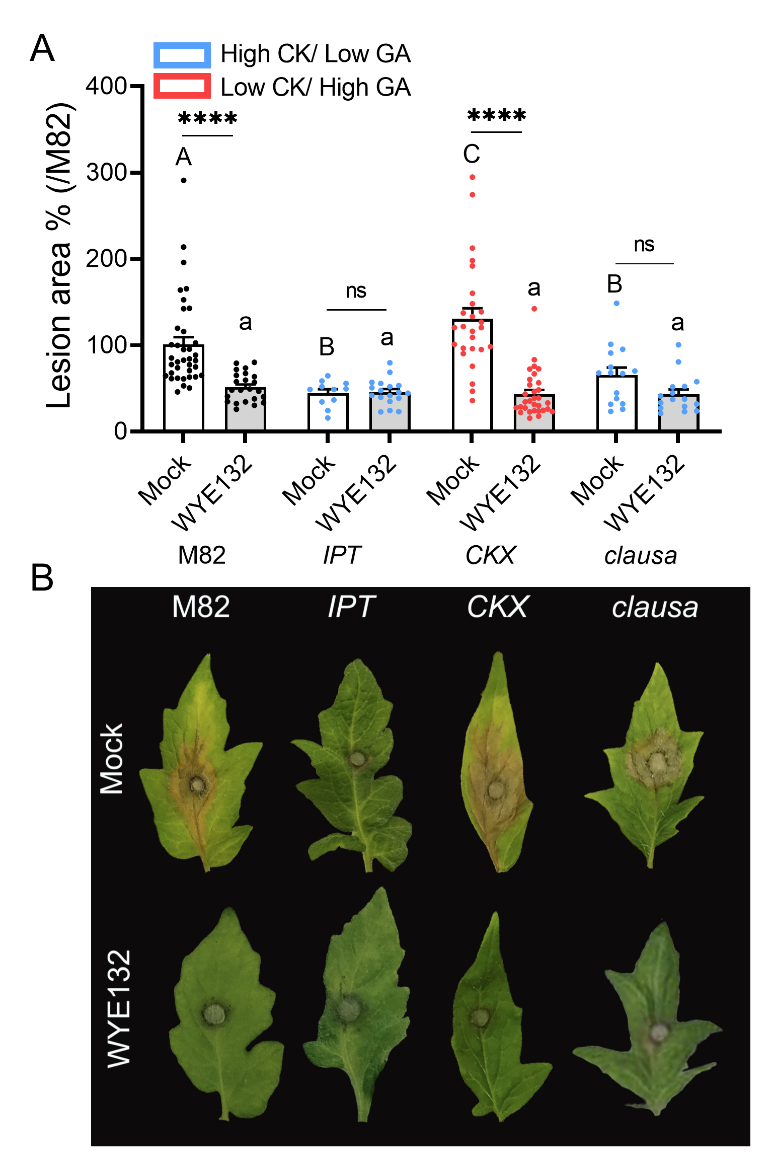

**Figure S1: TOR inhibition mediated disease resistance depends on the CK/GA balance- additional results**

*S. lycopersicum* plants of altered CK/GA genotypes: increased CK content *pBLS>>IPT7* ("*IPT*"), decreased CK content *pFIL>>CKX3* ("*CKX*"), and increased CK sensitivity and decreased GA sensitivity *clausa* mutant ("*clau*"), and their WT background M82, were treated with Mock (1:5000 DMSO in DDW) or 2 µM of the TOR inhibitor WYE132. Plants were challenged with *B. cinerea (Bc)* mycelia from a 72h old-culture, 24 h after drug treatment.

**A**: *Bc* necrotic lesion size. Experiment was repeated 4 independent times, bars represent mean ±SEM, all points shown. Asterisks indicate statistically significant disease reduction upon WYE132 treatment when compared with Mock treatment, in a one-way ANOVA with a Bonferroni post hoc test, N=20, ****p<0.0001, ns- non significant.

**B**: Representative images.

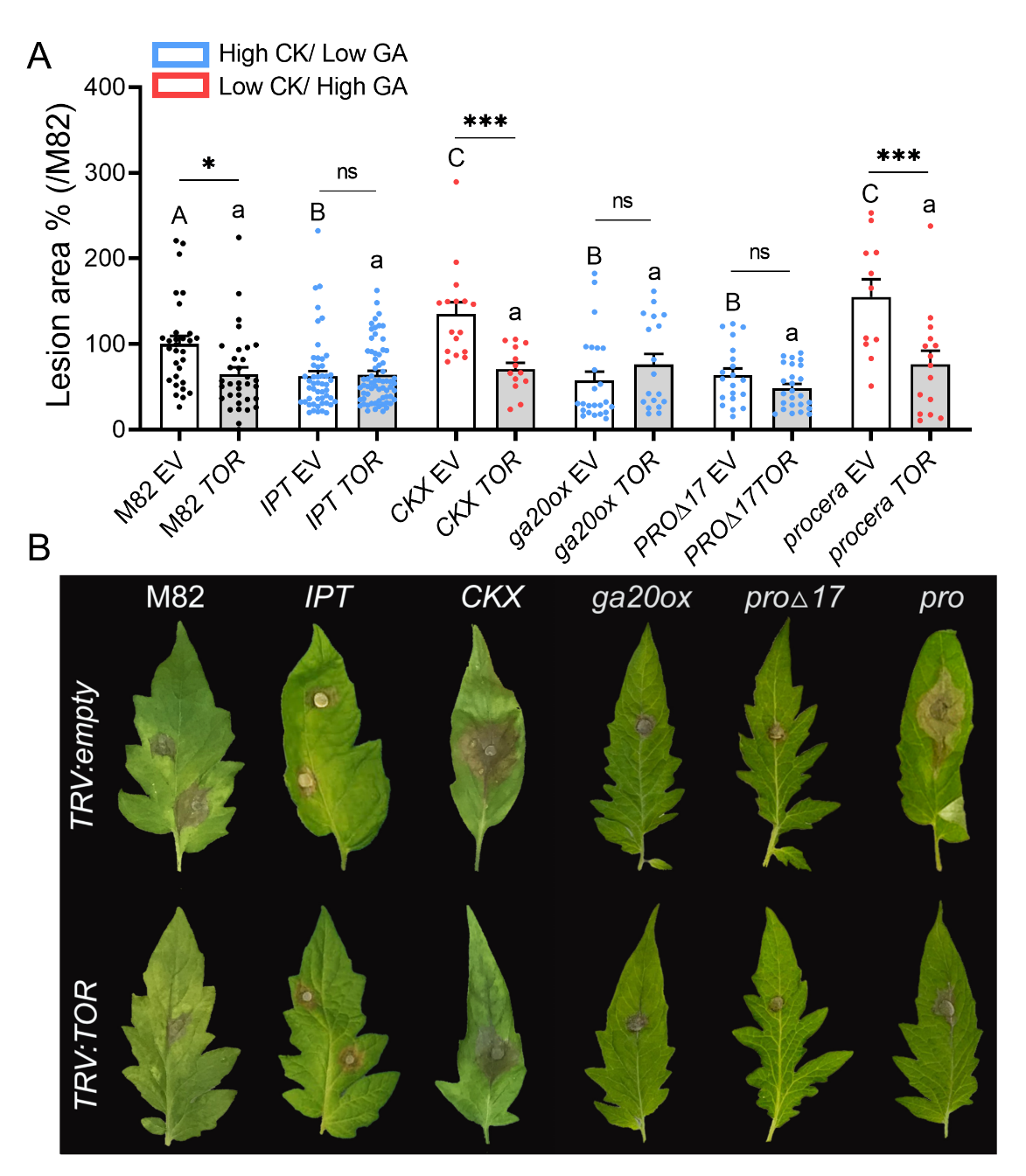

**Figure S2: *TOR* silencing mediated disease resistance depends on the CK/GA balance- VIGS results.**

*S. lycopersicum* plants of altered CK/GA genotypes: increased CK content *pBLS>>IPT7* ("*IPT*"), decreased CK content *pFIL>>CKX3* ("*CKX*"), decreased GA content mutant ("*ga20ox*"), decreased GA signaling *pFIL>>proΔ17* ("*proΔ17*"), increased GA signaling *procera* ("*pro*") and their WT background M82, were *TOR*-silenced using VIGS. 4 weeks after silencing, plants were challenged with *B. cinerea (Bc)* mycelia from a 72h old-culture.

**A**: *Bc* necrotic lesion size. Experiment was repeated 4 independent times, bars represent mean ±SEM, all points shown. Asterisks indicate statistically significant disease reduction upon *TOR* silencing when compared with control empty-vector silenced plants ("EV"), in a one-way ANOVA with a Bonferroni post hoc test, N>15, *p<0.05, ***p<0.001, ns- non significant.

**B**: Representative images.

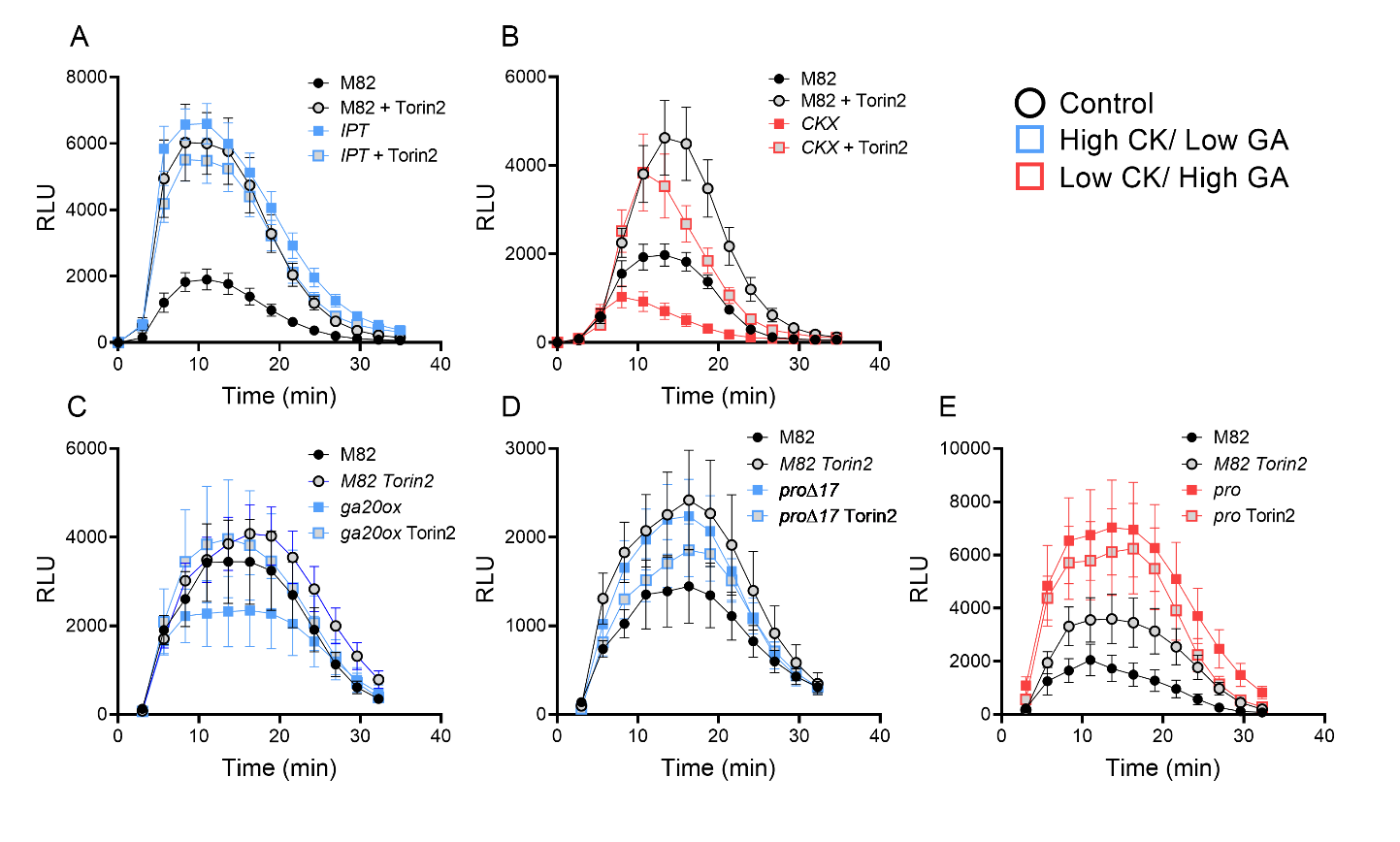

**Figure S3: TOR inhibition mediated disease resistance depends on the CK/GA balance- ROS kinetics**

*S. lycopersicum* plants of altered CK/GA genotypes: increased CK content *pBLS>>IPT7* ("*IPT*"), decreased CK content *pFIL>>CKX3* ("*CKX*"), decreased GA content mutant ("*ga20ox*"), decreased GA signaling *pFIL>>proΔ17* ("*proΔ17*"), increased GA signaling *procera* ("*pro*") and their WT background M82, were treated with Mock (1:5000 DMSO in DDW), or 2 µM Torin2. Plants were challenged with the immunity elicitor flg-22 (1 µM) 24 h after Torin2 treatment. ROS production was measured immediately after flg-22 application every three minutes, using the HRP-luminol method. Reaction Kinetics are plotted.

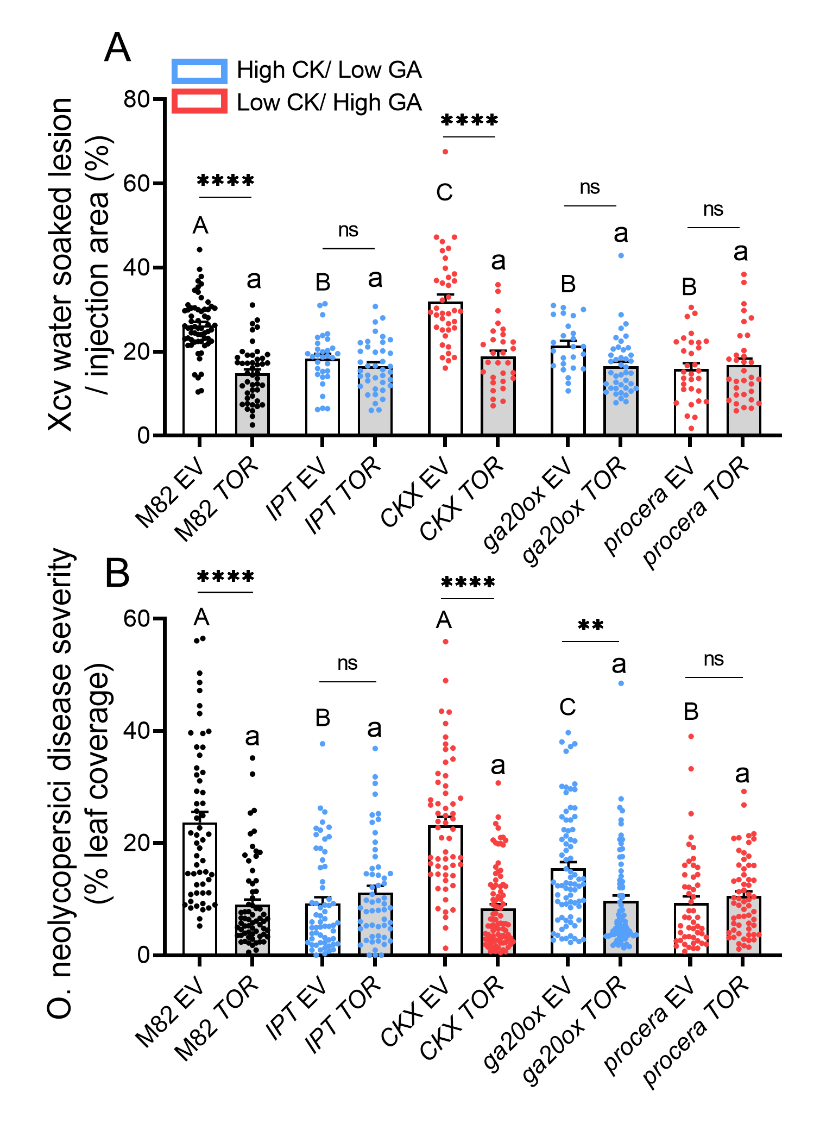

**Figure S4:** ***TOR* silencing mediated disease resistance to *X. campestris p. vesicatoria* and *O. neolycopersici* depends on the CK/GA balance**

*S. lycopersicum* plants of altered CK/GA genotypes: increased CK content *pBLS>>IPT7* ("*IPT*"), decreased CK content *pFIL>>CKX3* ("*CKX*"), decreased GA content mutant ("*ga20ox*"), and increased GA signaling *procera* ("*pro*") and their WT background M82, were *TOR*-silenced using VIGS. 4 weeks after silencing, plants were challenged with *X. camperstris p. vesicatoria* (A) or *O. neolycopersici* (B). Disease percentages were calculated after 10 (A) or 14 (B) days.

Bars represent mean ±SEM, all points shown. Experiments were repeated three independent times. Asterisks indicate statistically significant decreases in disease coverage upon *TOR* silencing. Different letters indicate statistically significant differences among samples, upper case letters for Mock silenced genotypes and lower case letters for *TOR*-silenced samples in Welch's ANOVA with Dunnett's post-hoc test. A: N>30, p<0.04. B: N>56, p<0.006.

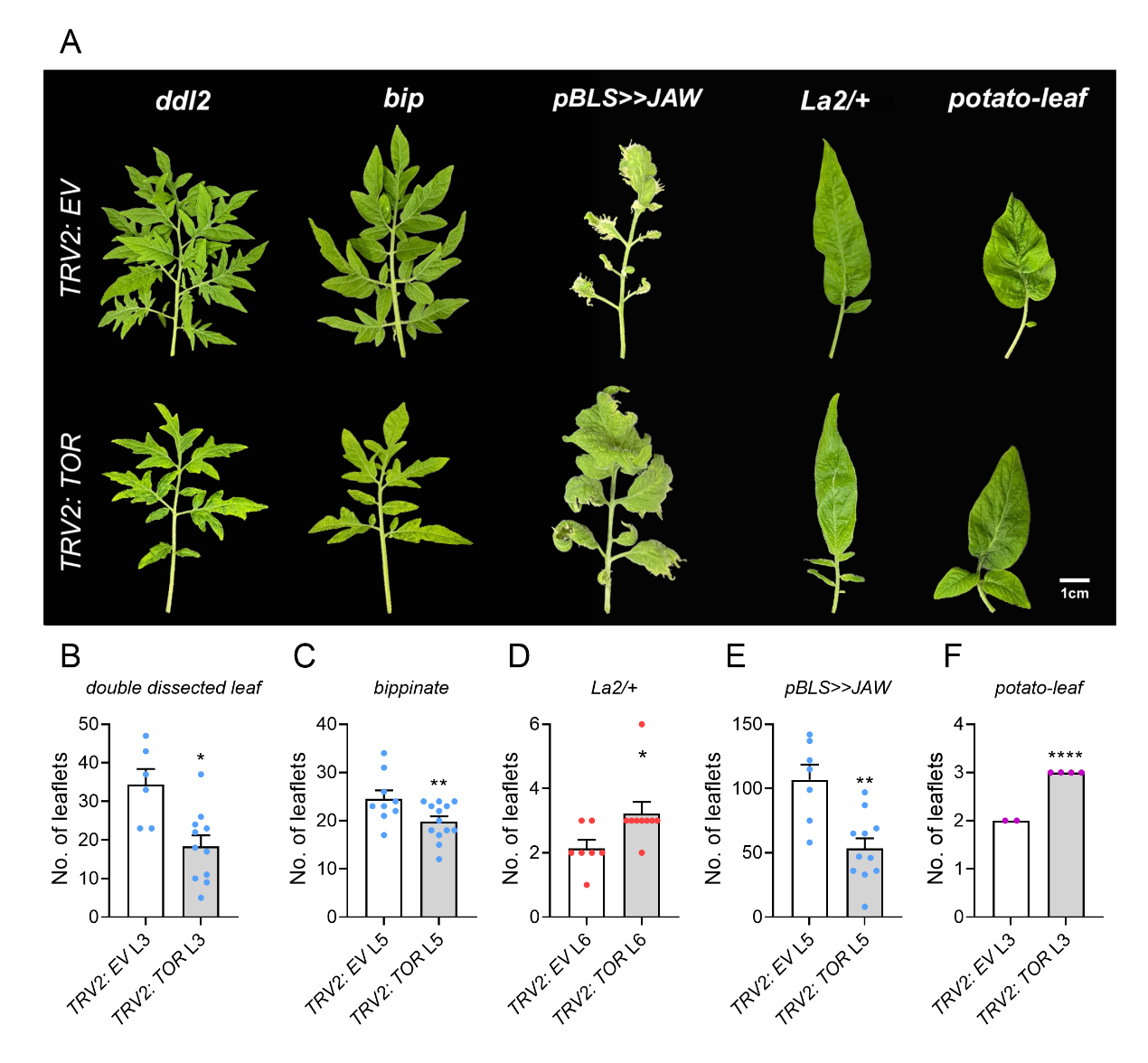

**Figure S5: TOR is required for the execution of developmental cues in the leaf**

*S. lycopersicum* plants mutated in the BELL transcription factors *Double Dissected Leaf*: *ddl2* (**A, B**), *Bippinate*: *bip* (**A, C**), the TCP transcription factor *Lanceolate*: *La2/+* (**A, D**), or overexpressing miR390 under the leaf specific promoter BLS: *pBLS>>JAW* (**A, E**), or the MYB transcription factor *C:* *potato leaf* (**A, F**), were *TOR*-silenced using VIGS. 4 weeks after silencing, leaf complexity was quantified by counting the leaflets on leaves 3, 5 or 6- as indicated.

Experiment was conducted 3 times. Boxplots represent inner quartile ranges (box), outer quartile ranges (whiskers), median (line in box), (B-E), or, bars represent Mean ±SEM (F), all points shown. Asterisks indicate statistically significant changes in leaf complexity upon *TOR* silencing as compared with the same leaf in empty-vector VIGSed plants, in a two-tailed t-test or a Mann-Whitney U test. **B:** N=6-11 individual plants, *p<0.05. **C:** N=9-14 individual plants, ***p<0.001. **D**: N=7-9 individual plants, **p<0.01. **E**: N=7-11 individual plants, *p<0.05. **F**: N=3-4 individual plants, ****p<0.0001.

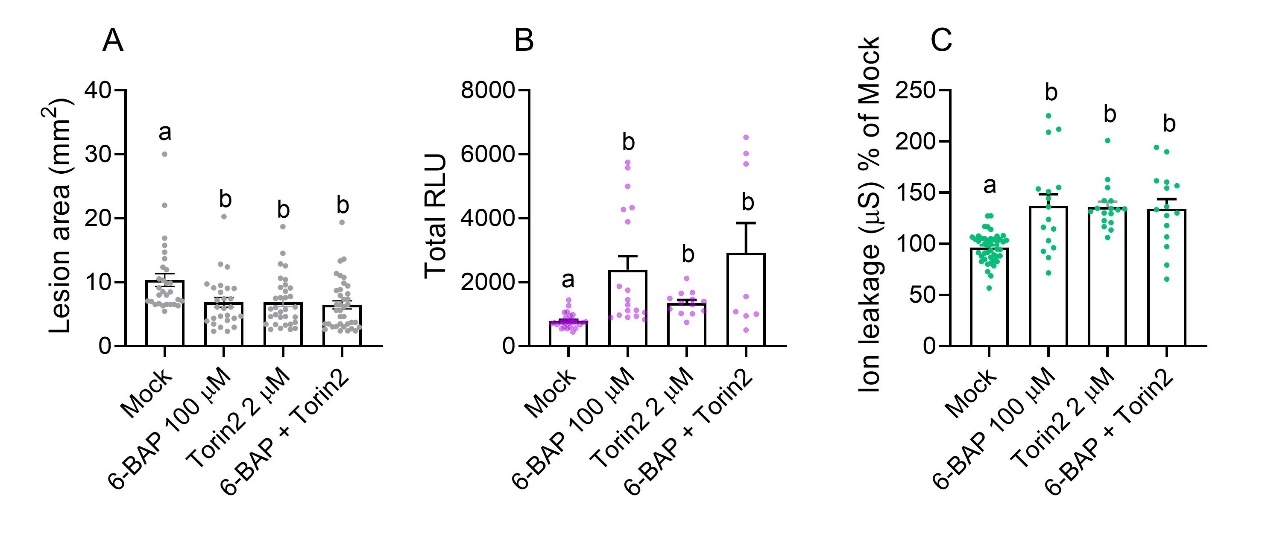

**Figure S6: TOR inhibition does not augment CK-mediated disease resistance.**

*S. lycopersicum* cv M82 plants were treated with 100 µM of the CK 6-benzylaminopurine (6-BAP), 2 µM Torin2, or a combination of both. Plants were challenged with *B. cinerea (Bc)* mycelia from a 72h old-culture (**A**), or with the immunity elicitors flg-22 (1 µM) (**B**) or Xylanase (1 µg mL^-1^)(**C**), 24 h after CK and Torin2 treatments. **A** Lesion area was quantified 72 h after *Bc* inocculation. Bars represent mean ±SEM, all points shown. Different letters indicate statistically significant differences in a one-way ANOVA with a Tukey post hoc test, N>25, p<0.017. **B** ROS production was measured immediately after flg-22 application every three minutes, using the HRP-luminol method, and expressed as the total Relative Luminescent Units (RLU) generated. Floating bars represent minimum to maximum values, line indicates median. Different letters indicate statistically significant differences in a one-way ANOVA with a Holm-Sidak post hoc test, N>8, p<0.01. **C** Conductivity of samples immersed in water for 40 h was measured. Mock average conductivity was defined as 100%. Bars represent mean ±SEM, all points shown. Different letters indicate statistically significant differences among samples in Welch's ANOVA with a Dunnett post hoc test, N>15, p<0.02.

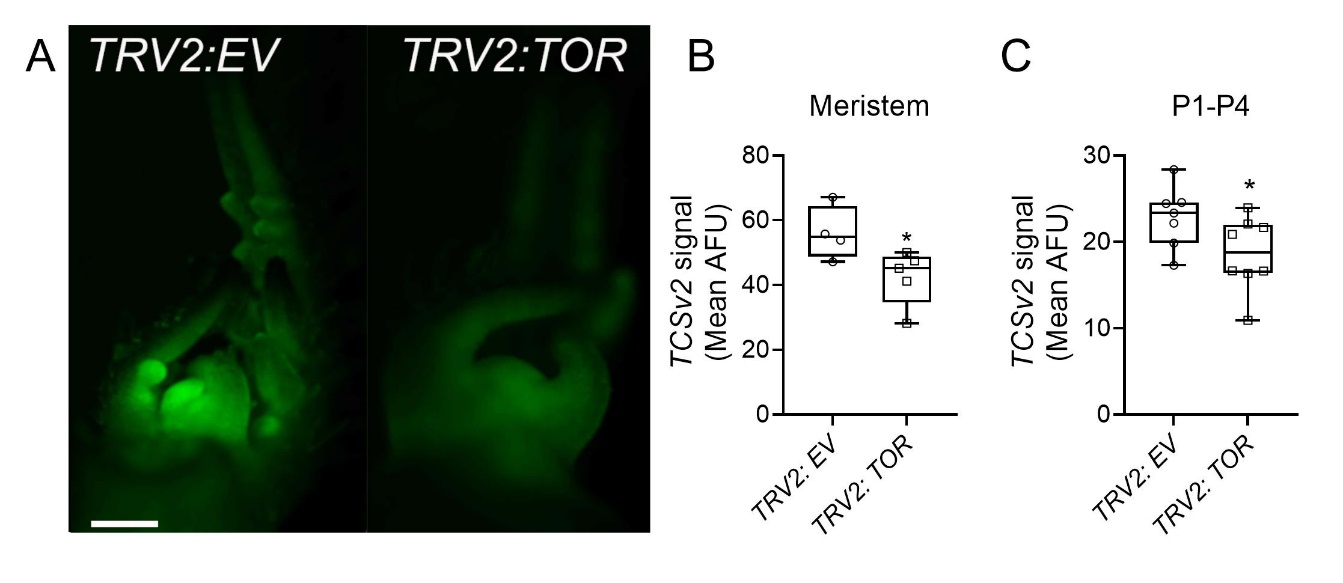

**Figure S7: *TOR* silencing by VIGS reduces CK response in the meristem and developing leaf primordia**

*S. lycopersicum* cv. M82 seedlings expressing VENUS driven by the cytokinin responsive promoter TCSv2 were *TOR*-silenced using VIGS. 2 weeks after silencing, TCSv2 driven total Venus fluorescence in the meristem (**A,B**) or youngest developing leaf primordia (**A, C**) was measured in images captured under identical conditions in shoots comprising the 4 youngest primordia. Boxplots represent inner quartile ranges (box), outer quartile ranges (whiskers), median (line in box), all points shown. Asterisks indicate significant TCSv2 signal reduction upon TOR silencing in a t-test, N>4, *p<0.05. Typical mock silenced and *TOR* silenced shoots are depicted in (**A**). Bar- 1000 µM.

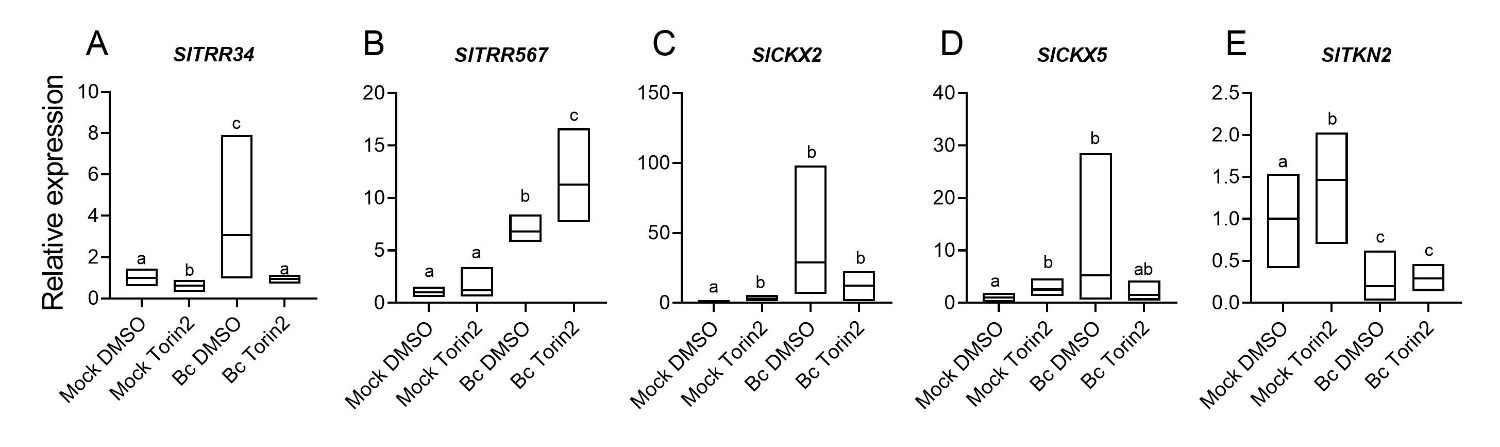

**Figure S8: TOR inhibition and *Bc* infection have differential effects on CK pathway genes**

Gene expression analysis of the indicated CK pathway and developmental genes, with and without Torin2 (2 µM) treatment, *Bc* infection, or combined *Bc* infection and Torin 2 treatment, was measured by RT-qPCR. Relative expression was calculated using the mean between the gene copy number obtained for three reference genes: RPL8 (Solyc10g006580), EXP (Solyc07g025390), and CYP (Solyc01g111170), and normalized to Mock treatment. Analysis was conducted on 6 biological samples comprised of 10 plants each, and repeated twice, N=12. Floating bars represent minimum to maximum values, line in bar represents the median. Different letters indicate statistically significant differences among samples in Welch's ANOVA (A, B), or Kruskal Wallis ANOVA (C, D) tests comparing each gene, A: p<0.013; B: p<0.047; C: p<0.001; D: p<0.008.

**Table S1: Primer pairs used in this work**

| **Locus** | **Name** | **Forward** | **Reverse** |
| --- | --- | --- | --- |
| Solyc01g080150 | *IPT3* | TTTCGCGGTGAAAAACTTCT | TGCAAGAAGGAAGTTGACGA |
| Solyc11g066960 | *IPT5* | CCAGTTGCAGCCAACAGAAT | GACCCCAGTTCTGCCCTCTA |
| Solyc12g014190 | *IPT6* | GATGTTCCAAAAGCCTCTCG | TAAACTTGCAAGCTCTGAGTCG |
| Solyc01g005680 | *LOG4* | GACAAAGGTGTGGAAGAAGGA | GCTCTTTGGGTGATGTAGCTG |
| Solyc08g062820 | *LOG5* | CATGTTGCTCCCCATGAAA | TGGAGACTGCTCCTTTGGAT |
| Solyc06g075090 | *LOG8* | TGAGCTTGGAAGAGAAATAGTATCAA | AAACCCATCAAACCAATGCT |
| Solyc05g006420 | *TRR3/4* | CGTCCCCTAAAGCATTCTCA | CGTCTTGTTGGTGATGTTGG |
| Solyc03g113720 | *TRR5/6/7* | GGGATTGATGGTTTGAAGGT | ATCTTGCTCAACACCGATGA |
| Solyc06g048600 | *TRR16B* | CATCAATGCATGGAAGAAGG | GCATTGCATTATTTGGCATC |
| Solyc01g088160 | *CKX2* | TGGTCCAAAATGGGATGTTT | AGGAGCAAGCATGGCTAAAG |
| Solyc04g016430 | *CKX5* | GAAGGATTTCGGCAACAGAT | GAAACCGATTTCGGATACAGA |
| Solyc12g008900 | *CKX6* | CAGGTGCTAAGCCATACTCTAGG | GGACATTCCATTAGGGGACA |
| Solyc07g066670 | *KS* | TGATTTCTTTGATGTAGGAGGTTC | GCTTGCCACTTAGATGCTTTG |
| Solyc04g083160 | *KO* | CCACGAAGACACGCAGGTAG | ATCGTTCAGGCTTCCACTCTT |
| Solyc01g080900 | *KAO* | CTTTCAAATCCAACAATCCTG | TTAAAACCTTCCTGCAACCT |
| Solyc03g006880 | *GA20ox1* | AGATTGTGTTGGTGGACTTCAA | TAGCGCCATAAATGTGTCG |
| Solyc06g035530 | *GA20ox2* | CGGTTTCTTTCTCGTGGCAA | TTTGCTTGTCGGAAAGTGGC |
| Solyc11g072310 | *GA20ox3* | ACTTTAGGGACAGGGCCTCA | ACTTGAAGCCCACCAACACT |
| Solyc03g119910 | *GA3ox2* | TTGGCCATGCATGCAAAACA | ATCTCGTCCCGTGTGTTTCC |
| Solyc02g089350 | *GAST1* | CAACAACAGAGAAATAACCAAC | TTATACGATGTCTTTGAACACC |
| Solyc11g011260 | *PROCERA* | TGATGCGACTATACTTGATATAAG | GGGTTAATCTGTTTAATAGAGTTC |
| Solyc07g061720 | *GA2ox4* | TTGAAAAGTTGGCGGAGGGA | AGCCTGAAAACAGAGTCGCT |
| Solyc07g061730 | *GA2ox5* | CAACACATCCGGCCTTCAAA | AACTCTGATCAGGTGGCACA |
| Solyc02g080120 | *GA2ox7* | AGCCACCTCCACTTCTCAAT | GGTTTGGCTGCTGTGACAAG |
| Solyc10g006580 | RPL8 | TGGAGGGCGTACTGAGAAAC | TCATAGCAACACCACGAACC |
| Solyc01g111170 | *CYP* | TGAGTGGCTCAACGGAAAGC | CCAACAGCCTCTGCCTTCTTA |
| Solyc07g025390 | *EXP* | TGGGTGTGCCTTTCTGAATG | GCTAAGAACGCTGGACCTAATG |
| Solyc11g005330 | *ACTIN* | TGGTCGGAATGGGACAGAAG | CTCAGTCAGGAGAACAGGGT |
| Solyc01g106770 | *TOR* (VIGS) | GGTCTAGAATGGCTGCCACCGTTCAGGCGATCCG | GGGGATCCTTCGCTGATGGTGACATCTAT |
| sg1-20ox-3 |  | taggtctcgTGCTTCTCACAAAGgttttagagctagaa | cgggtctctAGCAAAAAGCtgcaccagccggg |
| sg2-20ox-3 |  | taggtctcgAGTGGTTAACCATGgttttagagctagaa | cgggtctctCACTAAGAAAtgcaccagccggg |
| CRS-GA20ox3-1 |  | tgtggtctcaATTGCTTTTTGCTTCTCACAAAGgttttagagctagaaatagcaag |  |
| CRS-GA20ox3-2 |  | tgtggtctcaATTGTTCCACTAATAGACTTAAGgttttagagctagaaatagcaag |  |
